# LysoDCs in Peyer’s patches program compartmentalized microbiota-specific Th17 responses

**DOI:** 10.64898/2026.06.26.734008

**Authors:** Renan Oliveira Corrêa, Marie Cherrier, Rômulo G. A. Galvani, Rose Hardy, Antoine Robert, Marco M. Rodari, Cécilia Luciani, Mathieu Fallet, Bana Jabri, Nadine Cerf-Bensussan, Hugues Lelouard, Valérie Gaboriau-Routhiau

## Abstract

Segmented filamentous bacteria (SFB) are a canonical model of microbiota-driven Th17 immunity, but how distinct gut-associated lymphoid tissues shape the quality and effector potential of commensal-specific T-cell responses remains unclear. Here we show that Peyer’s patches (PPs) and mesenteric lymph nodes (MLNs) generate transcriptionally, clonally, and functionally distinct SFB-reactive CD4 T-cell programs. In PPs, CCR2-dependent monocyte-derived LysoDCs capture luminal SFB and locally prime antigen-specific CD4 T cells. PP priming drives robust T cell activation, Th17 differentiation with type 1 regulatory (Tr1)-like features, preferential clonal expansion within Th17-Tfh17 lineages, and tissue-retention programs. In contrast, CCR2-independent MLN priming induces a less differentiated, recirculating profile dominated by non-expanded clonotypes. Notably, these distinct programs carry functional consequences. Upon transfer into *Citrobacter rodentium*-infected lymphopenic mice, PP-primed T cells preserve barrier integrity and limit pathology, whereas MLN-primed cells from the same donors fail to provide equivalent protection. Together, these findings establish PP LysoDCs as specialized orchestrators of compartmentalized microbiota-specific immunity and identify the anatomical site of commensal priming as a key determinant of T-cell functional diversification and mucosal immune outcome.

**HIGHLIGHTS:** SFB colonization drives compartmentalized effector and regulatory CD4 T-cell programs in PPs versus MLNs.

Embigin⁺ LysoDCs in PPs directly sample SFB antigens and prime SFB-reactive CD4 T cells locally.

PP priming induces Tr1-like Th17 differentiation, preferential Th17-Tfh17 clonal expansion, and tissue residency.

PP-primed CD4 T cells ameliorate colitis-associated pathology during enteric infection, whereas MLN-primed cells do not.

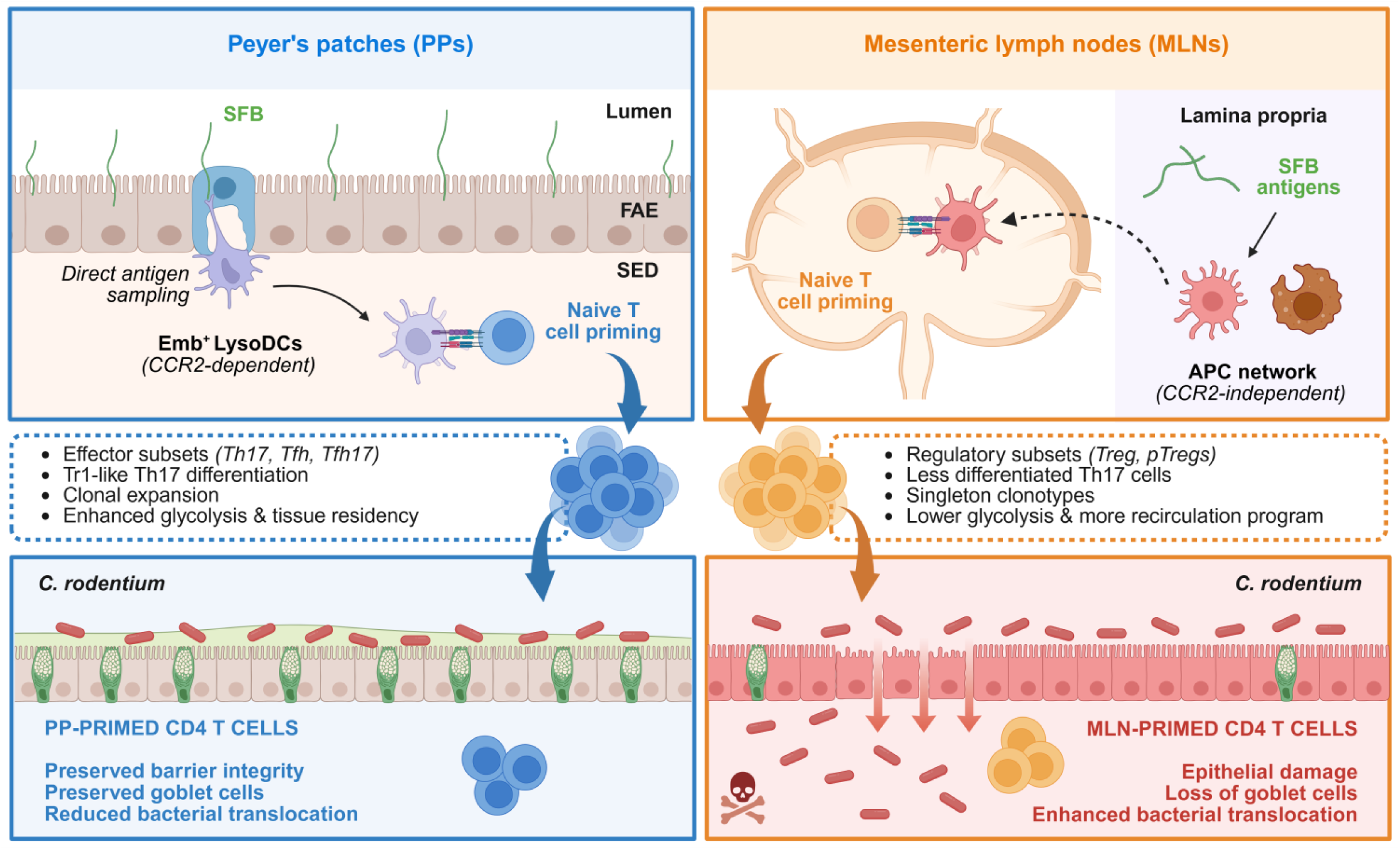

## INTRODUCTION

The intestinal mucosa is constantly exposed to a dense and diverse microbial community that shapes immune development, tissue homeostasis, and host defense ^1–3^. A central challenge in mucosal immunology is understanding how anatomically restricted antigen-presenting cell (APC) niches program distinct CD4 T-cell fates ^4^. This question is especially important for microbiota-driven T-cell responses, which must balance protective immunity with tolerance to avoid chronic inflammation ^5,6^. Segmented filamentous bacteria (SFB), which have recently been shown to be low-abundant but prevalent members of the human gut microbiota worldwide ^7^, provide a powerful model to address this issue. These commensals tightly adhere to the terminal ileal epithelium and are well known to elicit robust intestinal homeostatic Th17 and IgA responses ^8–10^ that contribute to protection against enteric pathogens ^11–16^. Yet, the cellular and anatomical mechanisms that initiate SFB-specific CD4 T-cell priming remain incompletely understood. In particular, although mesenteric lymph nodes (MLNs) have been proposed as the site of SFB-reactive Th17 induction ^17^, the responsible APCs and the relative contribution of other gut-associated lymphoid tissues, such as Peyer’s patches (PPs), have not been fully defined ^18–22^.

PPs and MLNs, the two main gut-associated lymphoid tissues, provide distinct immune environments that could drive divergent T-cell differentiation in response to the same microbial stimulus. PPs are specialized for sampling luminal antigens through the follicle-associated epithelium (FAE) and contain a unique network of APCs positioned to intercept microbial material arriving from the gut lumen ^23,24^. Among these, monocyte-derived LysoDCs occupy the subepithelial dome (SED), where they are strategically positioned to capture antigens delivered by M cells, as described for pathogenic bacteria such as *Salmonella* Typhimurium and *Listeria monocytogenes* ^23,25,26^. These cells are also capable of migrating to the interfollicular regions (IFRs), where they engage pathogen-reactive naive CD4 T cells and promote effector differentiation ^26,27^. Notably, sensing of mucosal-associated commensal bacteria by LysoDCs and conventional dendritic cells (cDCs) from PPs has been shown to induce Mincle-Syk axis-dependent stimulation of IL-17 and IL-22 responses *ex vivo*, as well as the development of Th17 cells *in vivo* ^28^. However, whether LysoDCs actively sample luminal commensal organisms *in vivo* and can directly prime antigen-specific T cells within PPs has not been demonstrated. Moreover, whether the same commensal bacterium, such as SFB, elicits qualitatively distinct CD4 T-cell responses when priming occurs in PPs versus MLNs, and whether such differences have functional consequences for mucosal immunity, remains unresolved.

Here, we show that SFB colonization induces compartmentalized CD4 T-cell priming in PPs and MLNs, with each site generating transcriptionally and functionally distinct responses. In PPs, we identify CCR2-dependent LysoDCs as specialized APCs that acquire luminal SFB and locally prime SFB-specific CD4 T cells, whereas SFB responses in MLNs occur in a CCR2-independent manner. Using single-cell transcriptomic and TCR repertoire analyses, we further show that PP- and MLN-primed cells diverge in activation state, migratory programs, differentiation trajectories, and clonal organization. Finally, we demonstrate that these differences have functional consequences *in vivo*, as PP-primed SFB-specific T cells better preserve mucosal integrity and limit pathology during enteric infection in lymphopenic hosts, whereas MLN-primed cells do not. Together, our findings establish PPs as a specialized site for early SFB-specific T-cell priming and identify LysoDCs as key orchestrators of compartmentalized microbiota-specific immunity. More broadly, these findings demonstrate that gut-associated lymphoid tissues are not merely distinct sites of priming for commensal-derived antigens, but that each site of priming triggers distinct functional programs in commensal-reactive CD4 T cells, with direct consequences for mucosal immunity.

## RESULTS

### SFB colonization induces compartmentalized Th17 responses in PPs and MLNs

To determine how PPs and MLNs contribute to SFB-specific CD4 T-cell priming, we first compared their capacity to support Th17 differentiation in a reductionist adoptive-transfer setting. Naive CD45.1 7B8 TCR-transgenic CD4 T cells ^29^ were transferred into CD45.2 ex-germ-free C57BL/6 adult mice one week after SFB monocolonization (Fig. S1A). Under these conditions, donor T cells underwent robust proliferation in both PPs and MLNs within 48 to 72 hours (Fig. S1B-C) and acquired comparable levels of RORγt expression (Fig. S1D-E). Importantly, no Th17 differentiation was observed when naive 7B8 T cells were transferred into SFB-negative SPF recipient mice (Fig. S1F-G), confirming SFB-specificity. These results indicate that both compartments can support SFB-dependent priming and Th17 differentiation when commensal antigens are broadly available.

We next asked whether differences in SFB priming would emerge under conditions in which antigen dissemination across tissues is not pre-established. To address this, we analyzed endogenous SFB-specific T cells following SFB monocolonization in adult germ-free mice by assessing the expression of the Vβ14 TCR chain, a widely used marker to track SFB-reactive CD4 T cells ^30^. At baseline, MLNs contained higher numbers of Vβ14^+^ T cells than PPs (Fig. 1A). Despite lower starting numbers, priming in PPs and MLNs initiated concurrently around day 3 post-colonization, peaking at day 7 with similar absolute numbers (∼4-6 × 10⁴ cells) (Fig. 1B). However, the subsequent kinetics diverged, with SFB-specific primed T cells declining in MLNs while remaining elevated in PPs (Fig. 1B). This was accompanied by a higher number of SFB-specific Th17 cells in PPs from day 7 onward, although slowly declining over time (Fig. 1C). In contrast, SFB-specific regulatory T cells (Tregs) were consistently low in PPs but accumulated progressively in MLNs (Fig. S1H), highlighting tissue-specific induction of distinct T-cell subsets by the same commensal. Thus, SFB colonization establishes a compartmentalized balance of effector and regulatory CD4 T-cell states across gut-associated lymphoid tissues.

**Figure 1.**
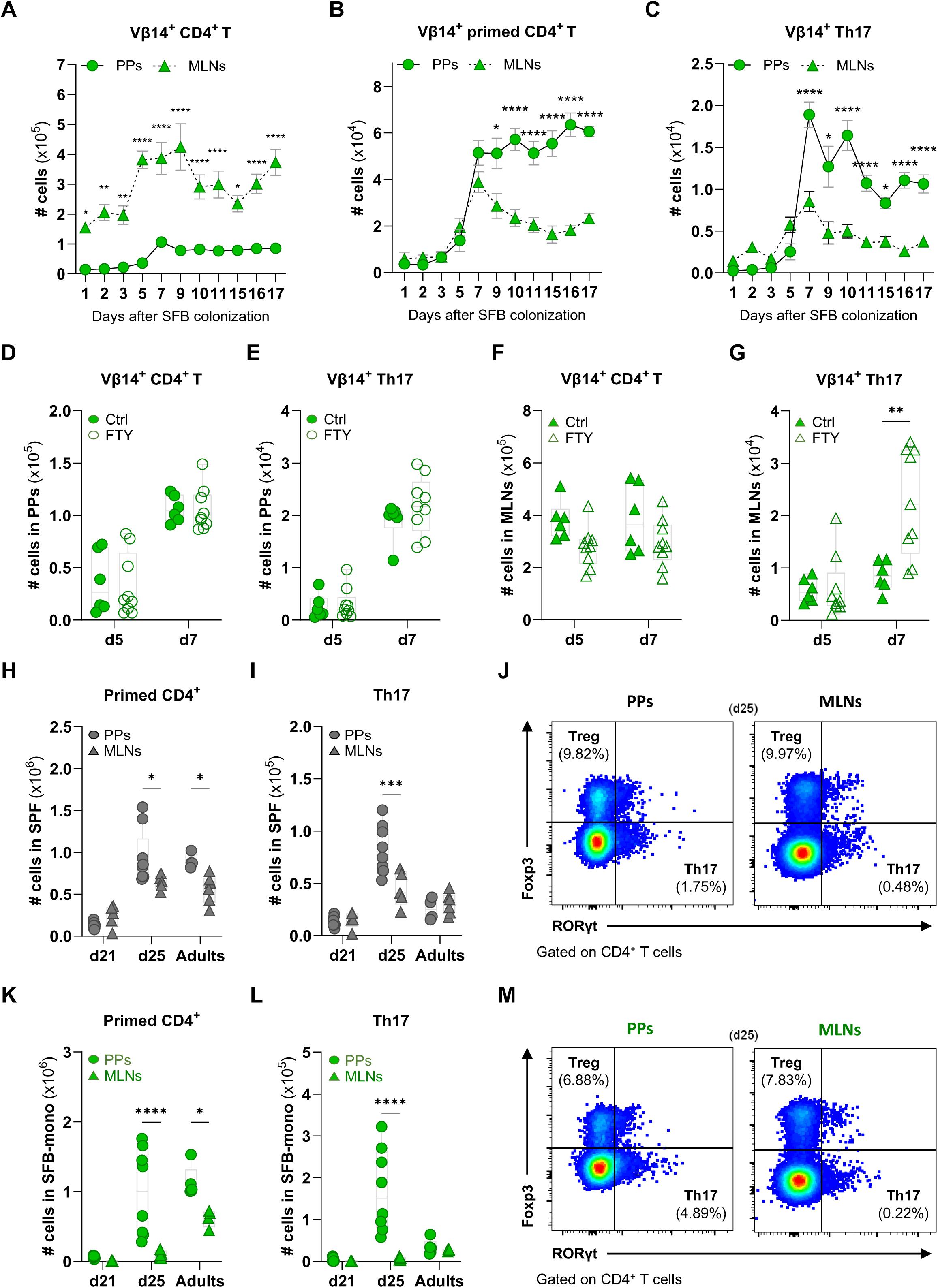
SFB colonization induces compartmentalized Th17 responses in PPs and MLNs (A-C) Absolute numbers of SFB-reactive Vβ14^+^ CD4 T cells (A), primed (CD44^hi^CD62L^-^) T cells (B), and Th17 (Foxp3^-^RORγt^+^) cells (C) in PPs (circles) and MLNs (triangles) of adult germ-free mice post-SFB monocolonization. Each symbol represents mean ± SEM. Two independent experiments, n = 4-10 mice/group/timepoint. (D-E) Absolute numbers of total CD4 T cells (D) and SFB-reactive Th17 cells (E) in PPs of mice treated with FTY720 (empty symbols) or vehicle (filled symbols), at days 5 and 7 post-SFB colonization. Data are presented as box-and-whisker plots with individual mice shown. Two independent experiments, n = 6-9 mice per group. (F-G) Same analysis in MLNs. (H-I) Absolute numbers of primed CD4 T cells (H) and Th17 cells (I) in PPs (circles) and MLNs (triangles) in SPF mice at weaning (postnatal d21), post-weaning (d25), and adulthood (d45-60). Data are pooled from 2-4 independent experiments (mean ± SEM). (J) Representative Foxp3/RORγt FACS plots, gated on live CD4^+^ T lymphocytes, from PPs and MLNs of SPF mice on postnatal d25. (K-M) Same analysis in SFB-monoassociated mice. (A-I, K-L) Statistical analysis by two-way ANOVA with Sidak’s multiple-comparisons test. See also Figure S1.

To determine whether these differences reflected distinct patterns of lymphocyte retention or egress, we treated ex-germ-free mice daily with the sphingosine-1-phosphate receptor antagonist FTY720, starting at the time of SFB monocolonization. In PPs, FTY720 treatment had little effect on the abundance of either total SFB-specific T cells or Th17 cells at days 5 and 7 (Fig. 1D-E, Fig. S1I), indicating a natural local retention of newly primed cells. In contrast, in MLNs, although total SFB-reactive T-cell numbers remained unchanged (Fig. 1F), Th17 cells accumulated by day 7 following FTY720 treatment (Fig. 1G, Fig. S1I), suggesting that newly polarized Th17 cells rapidly exit the MLNs under normal conditions. These data indicate that during the first week post-colonization, SFB-specific T cells follow distinct trafficking programs associated with their site of priming.

We then examined whether this compartmentalization also emerges under more physiological colonization settings. In SPF mice, SFB natural colonization begins around weaning time ^31,32^, peaks shortly thereafter, and then gradually declines, a dynamic mirroring the early-life colonization peak recently reported in humans ^7^. Interestingly, this kinetic was recapitulated in SFB-monoassociated mice vertically colonized from birth (Fig. S1J). During the early post-weaning period (postnatal day 25), SPF mice exhibited increased frequencies of primed CD4 T cells and Th17 cells compared to pre-weaning (postnatal day 21) mice, with these changes being more pronounced in PPs than in MLNs (Fig. 1H-J). Notably, this pattern was further accentuated in SFB-monoassociated mice (Fig. 1K-M). In adult mice, although PPs continued to harbor more primed CD4 T cells than MLNs, Th17 cell numbers declined and converged between tissues, suggesting that Th17 accumulation observed in PPs at early time points is not sustained over prolonged periods (Fig. 1H-M). This compartmentalization was also subset-specific, as Treg induction remained consistently lower in PPs but increased progressively in MLNs in both SPF and SFB-monoassociated mice (Fig. S1K-L).

Combined, these results demonstrate that SFB elicits a temporally dynamic but spatially biased T-cell response, with PPs favoring sustained local Th17 priming and tissue retention during early colonization, whereas MLNs support a more recirculating and regulatory-leaning response.

### LysoDCs internalize luminal SFB and engage SFB-specific T cells in interfollicular regions of PPs

The distinct SFB-specific CD4 T cell profiles observed in PPs and MLNs suggested that local APCs might provide specialized priming environments. We therefore investigated whether PP-specific LysoDCs, an enriched population in the SED of PP, can sample commensal SFB *in vivo* and interact with naive SFB-reactive CD4 T cells. In lysozyme M (LyzM)-GFP mice, in which LysoDCs are labeled ^25^, confocal imaging of ileal PPs showed LysoDC trans-M cell dendrites contacting luminal SFB at the FAE (Fig. S2A). Intraepithelial LysoDC dendrites also contained DNA-positive, dotted structures consistent with internalized SFB material, supporting direct luminal SFB uptake by LysoDCs (Fig. S2B).

Having established that LysoDCs can capture SFB, we next examined whether they can physically interact with SFB-specific T cells during priming. Naive CD45.1 7B8 CD4 T cells were transferred into CD45.2 LyzM-GFP recipient mice, and their ileal PPs were analyzed 72 hours post-transfer. Nearly all donor T cells within PPs localized to IFRs, where rare LysoDCs (GFP⁺MerTK⁺CD4⁻) formed contacts with both quiescent (Ki67⁻) and proliferating (Ki67^+^) SFB-specific T cells (Fig. 2A-B, Movie S1). Notably, LysoDCs preferentially associated with proliferating T cells, whereas LysoMacs (CD11c⁺MerTK⁺CD4⁺) and cDC2s (CD11c⁺MerTK⁻S100a4⁺) interacted with proliferating and non-proliferating cells at similar frequencies (Fig. 2C). Importantly, a subset of T cells contacting LysoDCs expressed RORγt but not Foxp3, consistent with newly primed Th17 cells (Fig. 2D, Movie S2).

**Figure 2.**
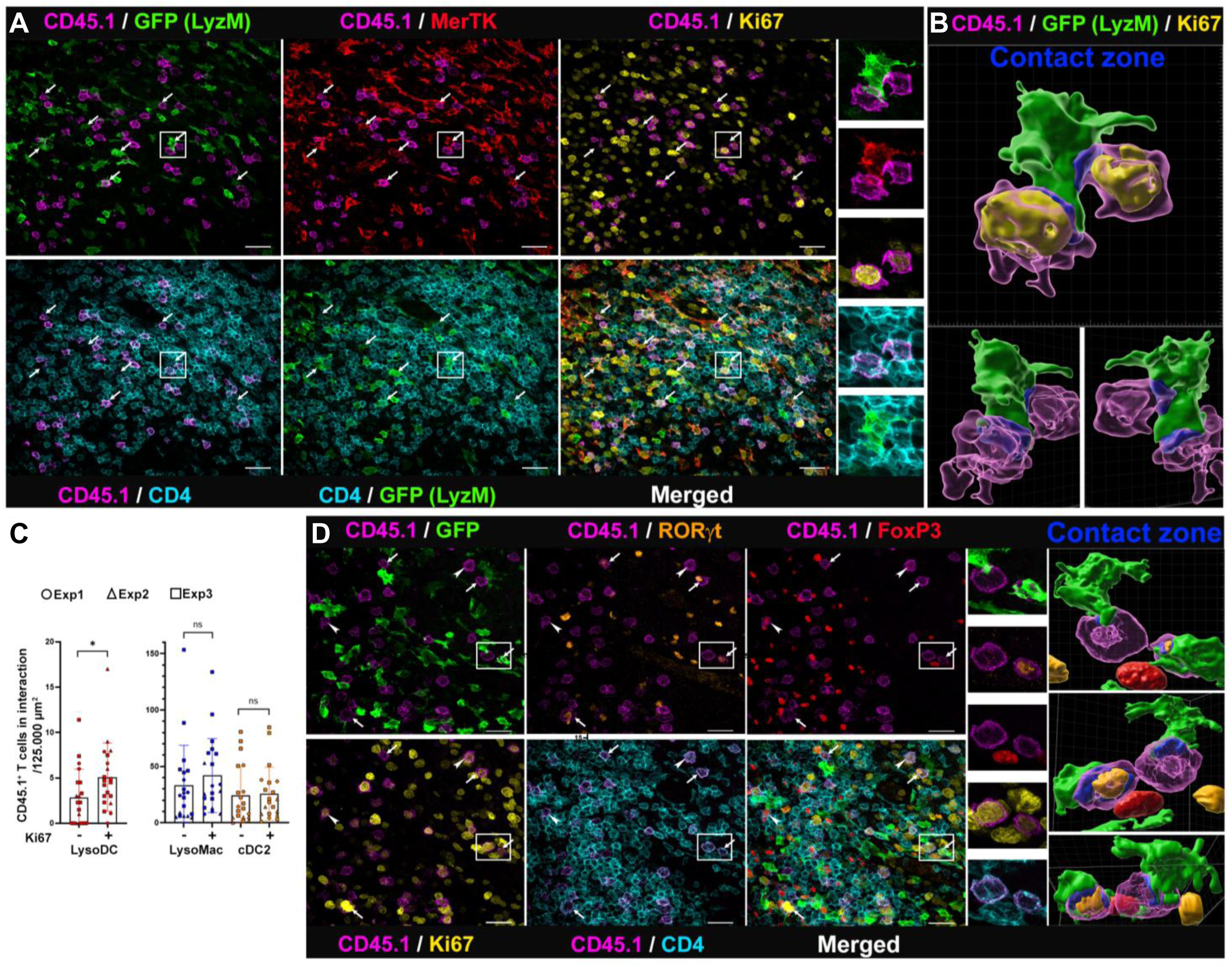
LysoDCs internalize luminal SFB and engage SFB-specific T cells in interfollicular regions of PPs Spectral confocal imaging of interfollicular region (IFR) of ileal PPs of CD45.2 LyzM-GFP recipient mice. (A) Transferred CD45.1 7B8 CD4 T cells (magenta), either quiescent or proliferative (Ki67^+^ in yellow), are observed in close interaction (arrows) with GFP+ (green) MerTK+ (red) CD4- (cyan) LysoDC. A higher magnification of the framed region is shown on the right. Scale bars, 20 µm. (B) 3D-reconstruction of a LysoDC (green) in close interaction (contact zone in blue) with two proliferating (Ki67^+^ in yellow) SFB-specific T cells (magenta). See Movie S1. (C) Quantification of interactions between LysoDCs (left graph) and LysoMacs or cDC2s (right graph) with quiescent (Ki67⁻) or proliferating (Ki67⁺) SFB-specific T cells. Three independent experiments, n = 3 mice/experiment. Statistical analysis by two-way ANOVA with Sidak’s multiple-comparisons test. ns = not significant. (D) A subset of proliferating (Ki67^+^ in yellow) SFB-specific T cells (magenta) in contact (arrows) with LysoDCs (green) expressed RORγt (orange) but not Foxp3 (red), indicating Th17 differentiation. Some others expressed only Foxp3 (arrowheads). A higher magnification of the framed region and a 3D reconstruction showing the contact zone (blue) between one LysoDC and a newly primed SFB-specific Th17 cell (RORγt^+^ nucleus in orange) are shown on the right. See Movie S2. See also Figure S2.

Together, these data indicate that despite their relative scarcity among interfollicular phagocytes, LysoDCs contribute to a specialized SFB-specific priming niche in PPs by sampling luminal bacteria and migrating to IFRs, where they directly engage naive SFB-reactive CD4 T cells.

### Embigin⁺ LysoDCs uniquely prime SFB-specific naive CD4 T cells *ex vivo*

To functionally confirm our imaging observations and determine whether LysoDCs can prime SFB-specific CD4 T cells, we isolated distinct APC subsets from PPs of adult ex-germ-free mice monocolonized with SFB for 14 days and assessed their ability to stimulate SFB-specific naive CD4 T cells *ex vivo*. To this end, sorted LysoDCs, cDC1s, and CD11b⁺ cDC2s were co-cultured with naive CD45.1 7B8 CD4 T cells either in the presence or absence of the SFB-derived A6 peptide ^29^ (Fig. 3A). Under peptide supplementation, all three APC subsets induced robust 7B8 T-cell proliferation, confirming their functional antigen presentation machinery (Fig. 3B-C). In contrast, when priming relied solely on endogenous SFB antigen acquired *in vivo*, only LysoDCs induced 7B8 T-cell proliferation, whereas cDC1s and CD11b⁺ cDC2s failed to do so (Fig. 3D-E). Because all three subsets presented exogenous peptide efficiently, this selective priming ability likely reflects the privileged access of LysoDCs to luminal SFB rather than differences in antigen presentation capacity. This priming activity was further restricted to SED-resident Embigin⁺ LysoDCs, as follicular-resident Embigin⁻ LysoDCs were also unable to induce T cell proliferation (Fig. 3D-E). Importantly, APCs isolated from PPs of SFB-negative SPF mice did not trigger T cell proliferation unless exogenous A6 peptide was supplemented, supporting the SFB specificity of our readout (Fig. S3A-B). Notably, under similar experimental conditions, APCs isolated from the MLNs or villus lamina propria of SFB-monoassociated mice also failed to trigger T cell proliferation in the absence of SFB peptide (Fig. S3C-D).

**Figure 3.**
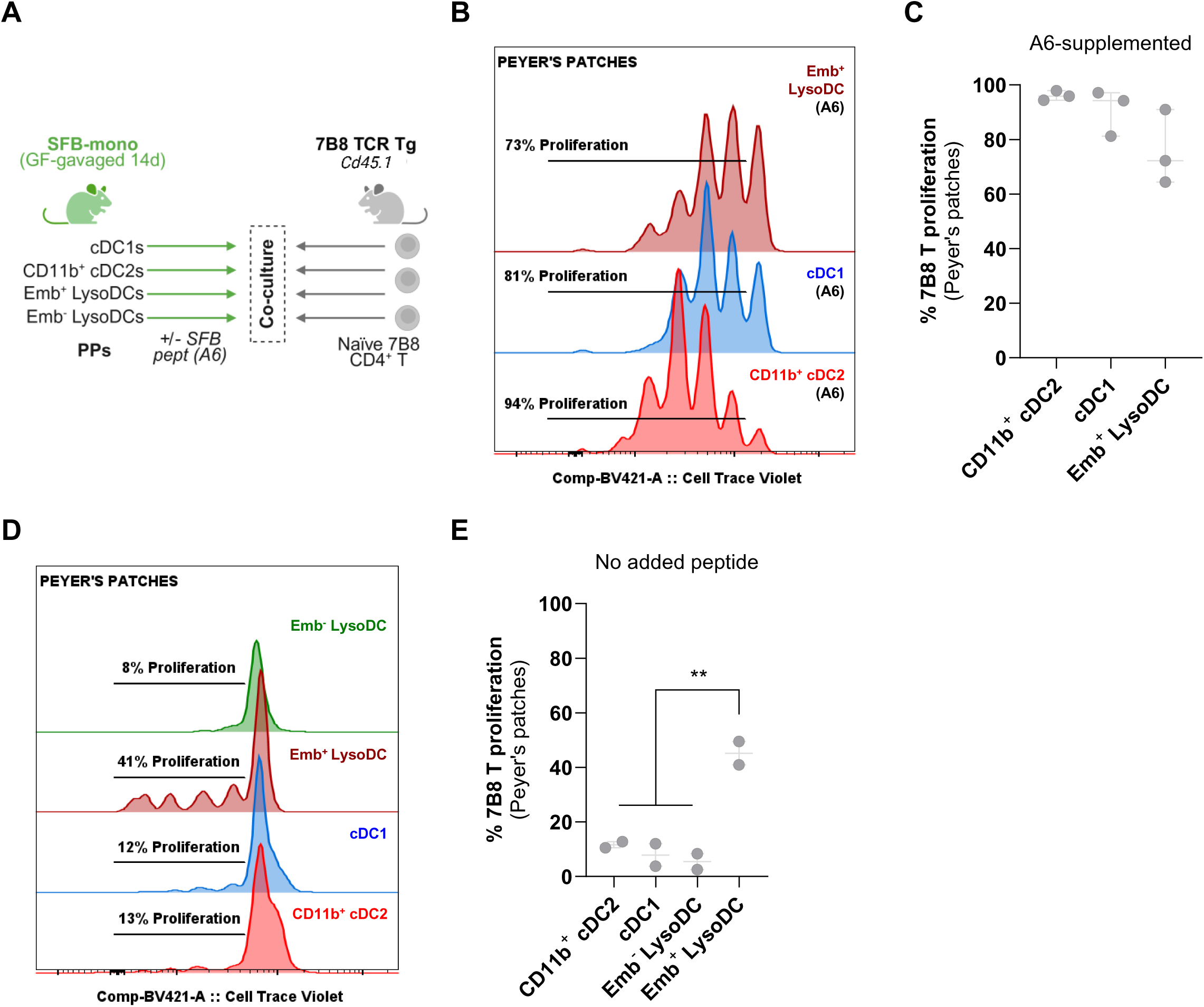
Embigin⁺ LysoDCs uniquely prime SFB-specific naive CD4 T cells *ex vivo* (A) Scheme of co-culture of distinct FACS-sorted APCs from PPs of SFB-monoassociated mice (ex-germ-free mice orally gavaged 14 days prior to the experiment) with naive 7B8 CD45.1 CD4 T cells, with or without SFB-specific A6 peptide. Cell proliferation was assessed by CTV dilution after four days in culture (related to B-E). (B-C) Representative FACS profiles (B) and frequencies of 7B8 T cell proliferation after co-culture with PP APCs in the presence of exogenous A6 peptide (C). Three independent experiments. In each experiment, APCs were FACS-sorted from pooled PPs of 15-20 mice, and 2-5 replicate co-cultures were established per condition. (D-E) Same co-culture analysis, but without exogenous A6 peptide supplementation. (C-E) Statistical analysis by one-way ANOVA with Sidak’s multiple-comparisons test. See also Figure S3.

Together, these results indicate that under steady-state conditions, Embigin⁺ LysoDCs in PPs are specialized APCs that efficiently capture native SFB antigens *in vivo*, process them efficiently, and present them in a form that supports the direct priming of SFB-specific CD4 T cells.

### CCR2-dependent LysoDCs are required for efficient SFB-specific priming in PPs

To further assess whether LysoDCs are required *in vivo* for the initiation of SFB-specific CD4 T-cell responses, we first used a clodronate-loaded liposome-based depletion strategy targeting their monocyte-derived precursors. Adult CD45.2 germ-free mice received a single dose of PBS- or clodronate-loaded liposomes intravenously. Five days later, mice were monocolonized with SFB by oral gavage combined with adoptive transfer of naive CD45.1 7B8 CD4 T cells, and were euthanized on day 10 post-injection (Fig. 4A). At this endpoint, persistent depletion of LysoDCs and maturing CD11b⁺ cDC2s was observed in PPs, whereas the levels of cDC1s, LysoMacs, and CD11b⁻ cDC2s had returned to baseline (Fig. 4B). This double depletion led to a pronounced reduction in Th17 differentiation of both donor-derived 7B8 T cells (Fig. 4C-D) and endogenous SFB-specific Vβ14 CD4 T cells (Fig. S4A-B) in the PPs. In the MLNs, although CD103⁻ cDC2s remained significantly reduced on day 10 post-clodronate injection (Fig. S4C), this depletion did not affect the induction of 7B8-derived or endogenous SFB-specific Th17 responses in this compartment (Fig. S4D-E).

**Figure 4.**
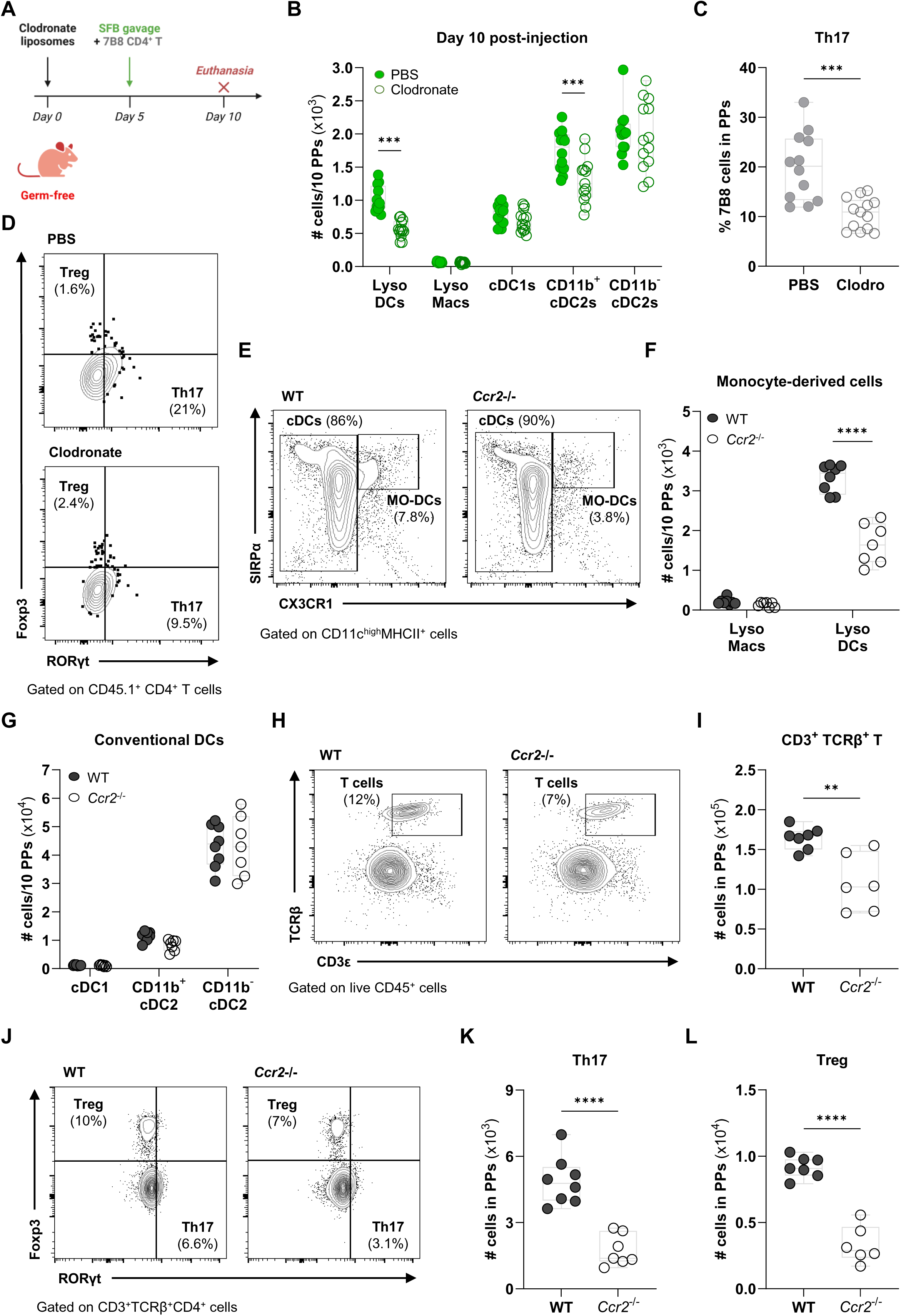
CCR2-dependent LysoDCs are required for efficient SFB-specific priming in PPs (A) Schematic of clodronate-loaded liposomes depletion treatment (related to B-D). (B) Number of mononuclear phagocytes per 10 PPs of clodronate-treated mice (empty symbols) or PBS-control mice (filled symbols) at day 10 post-liposome injection. Data show individual values for each mouse in box-and-whisker plots. Two independent experiments, n = 10-13 mice/group. Statistical analysis by two-way ANOVA with Sidak’s multiple-comparisons test. (C) Percentages of 7B8 Th17 (Foxp3^-^RORγt^+^) cells in PPs. Statistical analysis by unpaired two-tailed t-test. (D) Representative Foxp3/RORγt FACS plots, gated on live 7B8 (CD45.1^+^) CD4^+^ T lymphocytes, in PPs. (E-L) Comparisons between PPs of wild-type (WT, filled symbols) and *Ccr2^-/-^*mice (empty symbols). For quantified panels, data show individual values for each mouse in box-and-whisker plots. Two independent experiments, n = 6-8 mice/group. Statistical analysis by two-way ANOVA with Sidak’s multiple-comparisons test (F-G), or by unpaired two-tailed t-test (I, K-L). (E) Representative SIRPα/CX3CR1 FACS plots, gated on live CD11c^hi^MHCII^+^ cells. (F) Absolute numbers of monocyte-derived cells (LysoMacs and LysoDCs). (G) Absolute numbers of conventional dendritic cells (cDC1s, CD11b^+^ and CD11b^-^ cDC2s). (H) Representative TCRβ/CD3ε FACS plots, gated on live CD45^+^ cells. (I) Absolute number of CD3ε^+^ TCRβ^+^ T cells. (J) Representative Foxp3/RORγt FACS plots, gated on live CD3ε^+^ TCRβ^+^ CD4^+^ cells. (K-L) Absolute numbers of Th17 (Foxp3^-^RORγt^+^) (K) and Treg (Foxp3^+^RORγt^-^) cells (L). See also Figure S4.

Because clodronate treatment simultaneously depleted both LysoDCs and maturing CD11b⁺ cDC2s in PPs, this experiment alone could not distinguish the relative contribution of each population in the local induction of Th17 cells. We therefore turned to a more selective genetic approach using *Ccr2*-deficient mice, which specifically lack monocyte-derived LysoDCs in PPs while conventional DCs and resident macrophages are largely preserved ^27,33^. SPF wild-type (WT) and *Ccr2⁻^/^⁻* mice were orally gavaged with SFB at weaning (postnatal d21) and euthanized two weeks later. Quantitative PCR confirmed comparable SFB colonization between WT and *Ccr2⁻^/^⁻* mice (Fig. S4F), excluding differences in SFB load as a confounding factor. As expected, *Ccr2* deficiency caused a strong reduction of LysoDCs in PPs, with other major APC populations remaining largely unchanged (Fig. 4E-G). This selective depletion was associated with a reduction in total CD4 T cells, as well as diminished differentiation of Th17 and Treg in PPs (Fig. 4H-L). By contrast, although MLNs showed a similar decrease in monocyte-derived cells and unchanged conventional DC populations (Fig. S4G-I), no significant changes in total T-cell or Th17/Treg numbers were observed in MLNs under these conditions (Fig. S4J-M).

Together, these results demonstrate that SFB-specific T cell priming in PPs significantly relies on compartmentalized, CCR2-dependent recruitment of monocyte-derived LysoDCs. Whereas clodronate-based depletion highlights the overall contribution of monocyte-derived cells to this process, the *Ccr2⁻^/^⁻*model more precisely identifies LysoDCs as a non-redundant APC population required for the efficient generation of SFB-reactive T cell responses within the PP microenvironment.

### Compartment-specific priming instructs distinct transcriptional and clonal programs in SFB-reactive CD4 T cells

To dissect how PP and MLN priming environments shape SFB-specific CD4 T-cell fate, we performed single-cell RNA sequencing on FACS-sorted primed (CD44^hi^CD62L^-^) CD4⁺ T cells isolated from PPs and MLNs of adult SFB-monoassociated mice 7 days after colonization, corresponding to the peak of priming (Fig. 1B). Notably, because these mice were germ-free prior to SFB colonization, the primed CD4 T-cell pool is expected to be predominantly SFB-reactive. Individual mouse hashing enabled sample-level deconvolution.

Unsupervised clustering of comparable numbers of PP- and MLN-derived cells identified 19 populations including Th1, Th2, Th17, Tfh (encompassing GC Tfh, Tfh2, and Tfh17), Treg (resting tTregs and pTregs), and additional helper subsets, all represented in both compartments (Fig. 5A, S5A). Despite this shared diversity, PP- and MLN-derived cells differed markedly in composition. PPs were enriched in effector and follicular-associated populations, including Th17, Tfh17, GC Tfh, and cycling Tfh clusters, as well as exhausted Th2 and naive/Tcm IFN-primed cells. In contrast, MLNs were preferentially enriched in regulatory and migratory populations, including resting tTregs, pTregs, migratory effector-memory cells, Gilz-expressing naive/Tcm, and Th1 cells (Fig. 5B, S5B). These data indicate that SFB colonization establishes distinct compartment-specific CD4 T-cell programs that extend well beyond Th17 responses alone.

**Figure 5.**
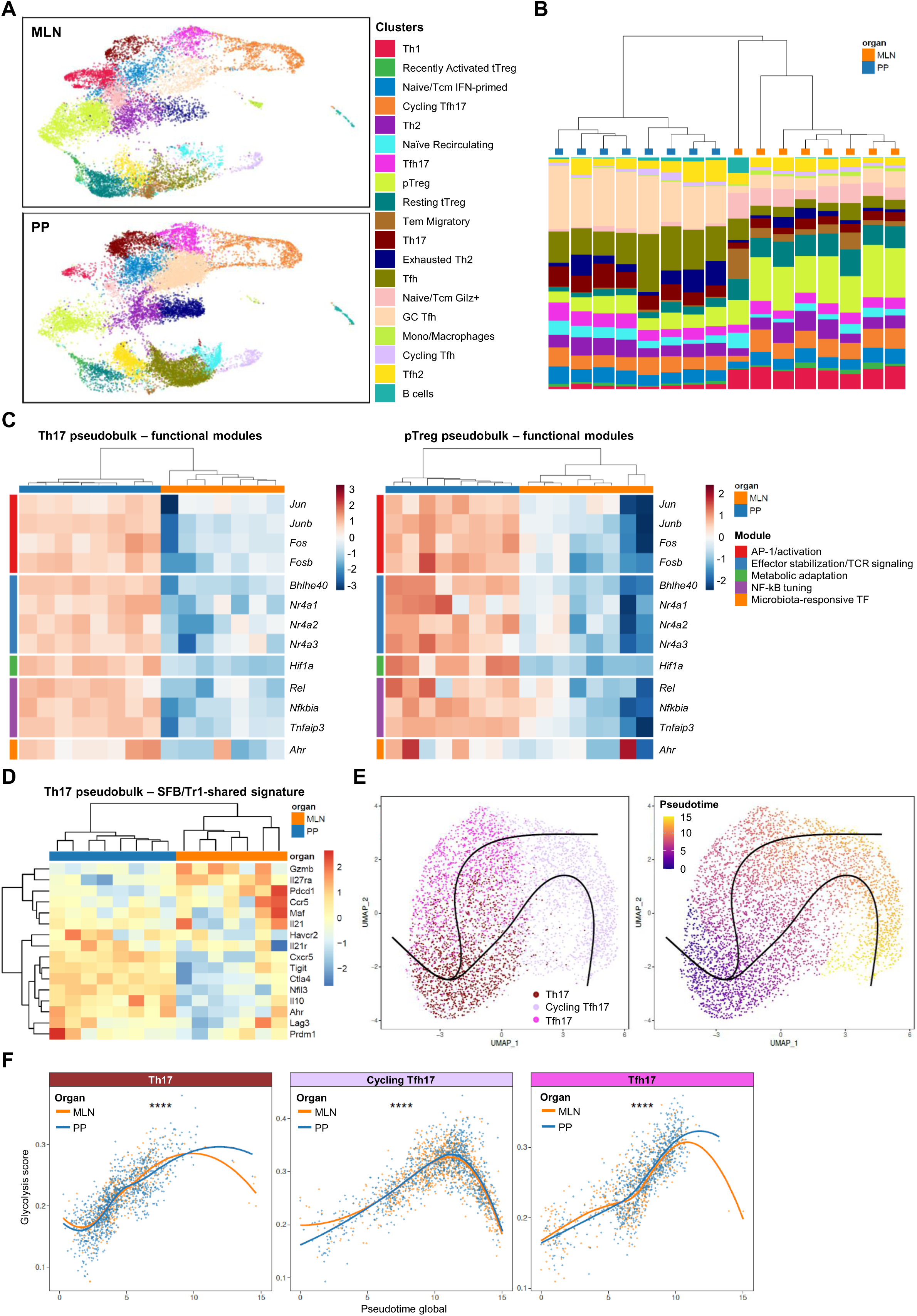
Compartment-specific priming instructs distinct transcriptional programming in SFB-specific CD4 T cells (A) UMAP visualization of primed (CD44^hi^CD62L^-^) CD4^+^ T cells from MLNs (top) and PPs (bottom) of ex-germ-free mice monoassociated with SFB for 7 days. Unsupervised clustering identified 19 populations, color-coded as indicated. Data represent 27,380 cells derived from PPs or MLNs, pooled from 8 mice, individually hashed for sample-level deconvolution. (B) Cluster composition by tissue origin. Each column represents one mouse; color indicates the relative proportion of each individual cluster (identified in A) within PP-derived (blue) and MLN-derived (orange) cells. Hierarchical clustering of columns groups tissues with similar composition. (C) Pseudobulk heatmaps showing expression of five selected functional gene modules in PP-(blue) and MLN-derived (orange) Th17 cells (left part) and pTreg cells (right part). Each column represents one mouse; color scale indicates z-scored expression. (D) Pseudobulk heatmap of SFB/Tr1-like shared signature gene expression in PP- (blue) and MLN-derived (orange) Th17 cells. Each column represents one mouse; color scale indicates z-scored expression. (E) UMAP visualization of Th17-related subsets (Th17, cycling Tfh17, and Tfh17) colored by cluster identity (left) or pseudotime value (right). Lines indicate the inferred differentiation trajectory from early Th17 states toward Tfh17 and cycling Tfh17 fates. (F) Glycolysis module scores plotted along global pseudotime for Th17 (left), cycling Tfh17 (middle), and Tfh17 (right) subsets. PP-derived cells (blue) and MLN-derived cells (orange) are shown separately. Lines represent loess-smoothed trends. Statistical analysis by Wilcoxon rank-sum test with Benjamini-Hochberg correction. See also Figure S5.

In addition to differences in cluster composition, gene expression analysis revealed that PP and MLN priming environments induce distinct transcriptional programs. PP-derived cells upregulated several coordinated gene-expression modules encompassing early activation/AP-1 signaling (*Jun*, *Junb*, *Fos*, *Fosb*), sustained TCR signaling (*Bhlhe40*, *Nr4a1/2/3*), metabolic adaptation (*Hif1a*), NF-κB regulation (*Rel*, *Nfkbia*, *Tnfaip3*), effector cytokine production (*Il17a*, *Il22*, *Il23r*), microbiota responsiveness (*Ahr*), and tissue residency (*ItgaE*, *Cd69*) (Fig. S5C, S5D). In contrast, MLN-derived counterparts preferentially expressed regulatory and recirculation-associated programs, including less differentiated cell markers (*Foxp3*, *Il2ra*, *Nrp1*, *Ctla4, Sell*) and S1P receptor-dependent egress programs (*S1pr1*, *Klf2*) (Fig. S5C, S5D). Notably, the PP-associated transcriptional signature was elevated across nearly all clusters in PPs compared to their MLN-derived counterparts (Fig. S5D) regardless of their tissue enrichment profile, as illustrated by both PP-enriched Th17 cells (Fig. 5C, left) and MLN-enriched pTregs (Fig. 5C, right). This indicates that the priming microenvironment, rather than cell lineage identity, acts as a dominant instructive cue shaping shared transcriptional programs across commensal-activated T helper subsets.

Considering that SFB-induced Th17 cells in the ileal lamina propria share a subset of signature genes with IL-10⁺Foxp3⁻ type 1 regulatory (Tr1) cells ^34^, we then used this shared gene set to compare PP- and MLN-primed Th17 cells. We found that many of these Tr1-associated genes were upregulated in PP-derived Th17 cells (Fig. 5D), suggesting that PP priming may initiate this Th17 regulatory program locally, whereas MLN-derived Th17 cells may only fully acquire these features upon reaching the ileal lamina propria ^34^. Additionally, pseudotime trajectory analysis of Th17-related subsets from PPs and MLNs revealed a continuum in which Th17 cells occupied early positions and progressed toward Tfh17 and/or cycling Tfh17 states (Fig. 5E), consistent with local plasticity toward Tfh-like differentiation. Along these trajectories, PP-derived cells exhibited increased glycolytic activity at both early (Th17) and more advanced (Tfh17 and cycling Tfh17) differentiation stages compared to their MLN-derived counterparts (Fig. 5F), indicating that the PP priming microenvironment promotes faster and sustained glycolysis-biased metabolic reprogramming during T-cell differentiation. These data reinforce the model in which the anatomical site of priming acts as a dominant instructive cue that coordinately imprints transcriptional, functional, differentiation, and metabolic programs across commensal-specific CD4 T-cell states.

To determine whether these compartmentalized transcriptional programs were accompanied by distinct clonal architectures, we next analyzed the TCR repertoire of PP- and MLN-derived primed CD4 T cells. PPs contained a higher number of unique TCRαβ clonotypes and showed limited overlap with MLNs (Fig. 6A). This pattern was observed despite broadly similar TRAV and TRBV gene usage across compartments (Fig. S6A-C), suggesting that it was not driven by differences in V-gene usage. Clonotype distribution further revealed pronounced tissue-specific segregation, with many clonotypes restricted to individual clusters within each tissue (Fig. 6B-C). MLNs were largely dominated by singleton clonotypes, whereas PPs contained a higher proportion of small and medium-sized expanded clones that were preferentially enriched in effector populations, particularly Th17 and cycling Tfh17 clusters (Fig. 6B, 6D-E). Shared clonotypes between cell types were most frequently observed among PP-derived Th17-like subsets, including Th17, cycling Tfh17, Tfh17, and occasional pTreg clusters (Fig. 6F), indicating that the shared repertoire is concentrated within PP effector-like lineages, consistent with the differentiation trajectories observed in pseudotime analysis.

**Figure 6.**
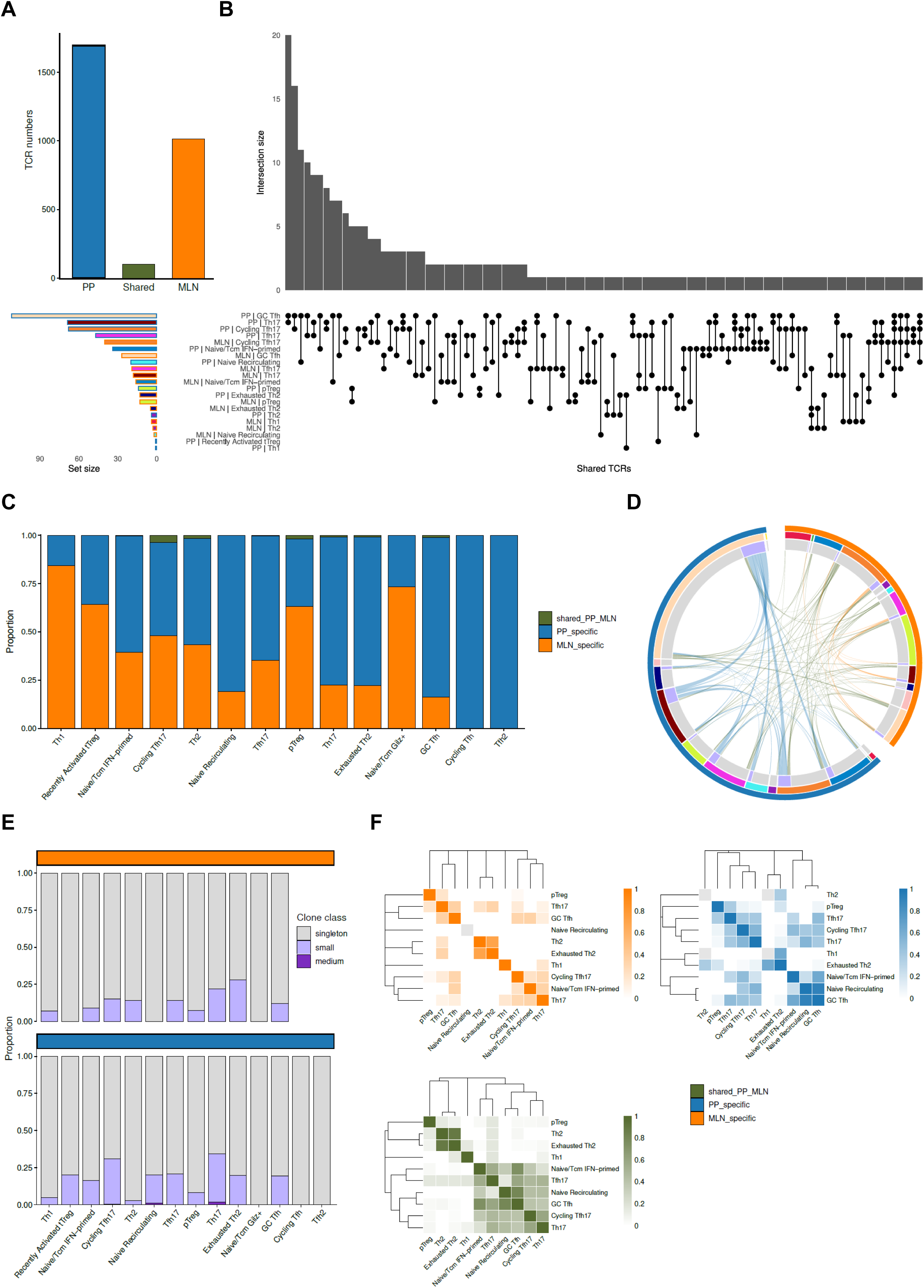
Clonal analysis reveals site-associated expansion of SFB-specific CD4 T cells (A) Total number of unique TCRáâ clonotypes identified in PPs (blue), MLNs (orange), and shared between both tissues (green). (B) UpSet plot showing clonotype sharing across tissue-cluster combinations. Each column represents a unique intersection of cluster-tissue pairs sharing clonotypes. Rows indicate the tissue-cluster identity, and connected dots mark the groups sharing a given set of clonotypes. Bar height indicates the number of shared clonotypes per intersection. Horizontal bars (bottom left) indicate the total number of clonotypes per tissue-cluster combination. (C) Proportion of clonotypes per cluster classified as PP-specific (blue), MLN-specific (orange), or shared between tissues (green). (D) Chord diagrams showing clonotype sharing between clusters within PPs (left pair, blue) and MLNs (right pair, orange). Arc width is proportional to the number of shared clonotypes between connected clusters. (E) Clonal expansion analysis within each cluster, shown separately for MLN (top) and PP (bottom). Proportion of clonotypes classified as singletons (gray), small clones (light blue), or medium clones (dark blue). (F) Clonotype transition probability heatmaps showing the likelihood that a clonotype found in one cluster is also found in another cluster, calculated separately for MLN (top left, orange), PP (top right, blue), and shared clonotypes (bottom, green). Color intensity indicates transition probability. See also Figure S6.

Collectively, these data indicate that the anatomical site of priming shapes both transcriptional programs and clonal architecture of SFB-specific CD4 T cells. PPs support preferential clonal expansion and diversification within effector-like subsets, whereas MLNs are characterized by a predominance of non-expanded, regulatory, and recirculating clonotypes.

### PP-primed SFB-specific T cells confer enhanced mucosal protection during *Citrobacter rodentium* infection

Finally, to test whether these compartment-specific priming programs result in distinct functional outcomes in the intestinal mucosa, we compared the capacity of PP- and MLN-primed SFB-specific CD4 T cells to provide protection against enteric infection *in vivo*. Germ-free *Rag1⁻^/^⁻* mice were orally gavaged with 2×10⁹ CFUs of *Citrobacter rodentium* (CR), an attaching-effacing pathogen whose clearance partially depends on adaptive immunity ^35,36^ and for which SFB colonization has been reported to confer protection ^9^. Twenty-four hours post-infection, these infected mice received equal numbers of FACS-sorted primed (CD44^hi^CD62L^-^) CD4 T cells (4x10^5^ cells/mouse) isolated from PPs or MLNs of SFB-monoassociated donor mice that had been colonized for 7 days. Importantly, donor mice were treated daily with FTY720 throughout the 7-day colonization period to block lymphocyte recirculation and ensure that sorted cells reflect tissue-restricted priming programs. Control groups included CR-infected mice receiving PBS (no T cells) and non-infected *Rag1⁻^/^⁻* germ-free mice. In lymphopenic recipients, transferred T cells are expected to undergo homeostatic proliferation ^37,38^, enabling their intrinsic functional programs to manifest *in vivo*.

We observed that all CR-infected mice showed comparable initial weight loss up to day 13 post-infection. However, from day 16 onward, mice receiving PBS alone or MLN-primed cells developed accelerated and severe weight loss (>20% by day 20), whereas those receiving PP-primed cells maintained significantly milder weight loss (∼10% at termination) (Fig. 7A). Weight of non-infected control mice remained stable throughout the experiment (Fig. 7A, dotted line). Consistent with reduced disease severity, colon shortening (a hallmark of colitis) was less pronounced in PP-cell recipients compared to PBS- and MLN-cell groups (Fig. 7B). Fecal CR burdens remained similar across groups on the day prior to euthanasia (Fig. S7A), indicating that the differences in disease outcome were not attributable to differences in pathogen colonization dynamics.

**Figure 7.**
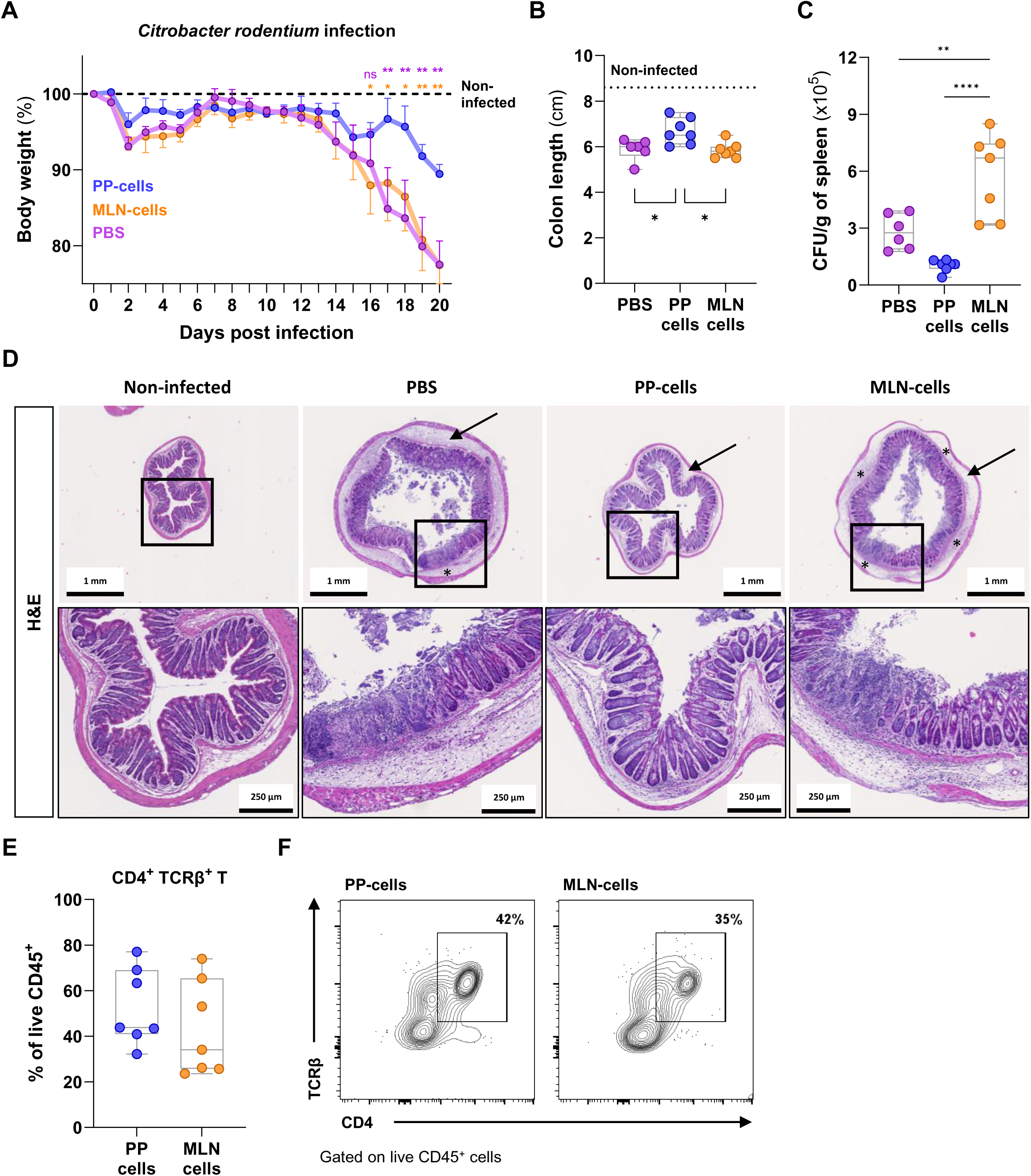
PP-primed SFB-specific T cells confer enhanced mucosal protection during Citrobacter rodentium infection (A) Body weight loss relative to initial weight in *Citrobacter rodentium* (CR)-infected *Rag1*-deficient mice receiving PBS only (purple) or primed CD4 T cells from PPs (blue) or MLNs (orange) of SFB-monoassociated mice treated with FTY720 throughout colonization. The dotted line indicates the mean from non-infected germ-free *Rag1*-deficient mice. Data are pooled from two independent experiments and shown as mean ± SEM. n = 4-8 mice per group. Statistical analysis by two-way ANOVA with Sidak’s multiple-comparisons test. (B) Colon length (cm) at the time of euthanasia. The dotted line indicates the mean colon length from non-infected germ-free *Rag1*-deficient mice. Data show individual values in box-and-whisker plots. (C) CR bacterial burden in the spleen, quantified by culture and CFU counting, and normalized to spleen weight. Data show individual values in box-and-whisker plots. (B-C) Statistical analysis by one-way ANOVA with Tukey’s multiple-comparisons test. (D) Representative H&E-stained distal colon sections from *Rag1*-deficient mice, either non-infected or infected with CR and receiving PBS or the indicated primed cells. Black arrows indicate submucosal edema; black asterisks indicate epithelial ulceration. Scale bars: 1 mm (top) and 250 µm (bottom). (E) Percentage of CD4⁺TCRβ⁺ T cells among live CD45⁺ cells isolated from the colonic lamina propria of mice transferred with PP- or MLN-primed CD4 T cells. Data show individual values in box-and-whisker plots. Statistical analysis by unpaired two-tailed t-test. (F) Representative CD4/TCRβ FACS plots, gated on live CD45⁺ cells. See also Figure S7.

Analysis of CR translocation to systemic sites showed markedly increased bacterial dissemination to the liver (Fig. S7B) and spleen (Fig. 7C) in PBS- and MLN-cell recipients compared to PP-cell recipients, indicating enhanced intestinal barrier protection conferred by transferred PP-primed T cells. Histological analysis of the distal colon further supported this conclusion. Despite comparable crypt hyperplasia across all CR-infected groups compared to non-infected mice (Fig. S7C), PP-cell recipients showed preserved goblet cell numbers (Fig. S7D), better preserved epithelial architecture with reduced submucosal edema (black arrows), and limited immune cell infiltration (Fig. 7D). In contrast, MLN-cell recipients exhibited severe epithelial damage and ulceration (black asterisks) with pronounced inflammatory infiltrates, comparable to PBS-treated controls (Fig. 7D). Importantly, FACS analysis of transferred CD4 T cells recovered from the colonic lamina propria of infected *Rag1⁻^/^⁻* recipients at the time of euthanasia revealed comparable levels of CD4 T-cell reconstitution (Fig. 7E-F) and equivalent frequencies of RORγt⁺ Th17 cells (Fig. S7E-F) in both groups, ruling out differences in T cell engraftment or Th17 representation as confounding factors.

Collectively, these results indicate that the anatomical site of commensal-induced CD4 T-cell priming influences their subsequent effector function, with SFB-specific PP-primed cells conferring superior protection against *C. rodentium* infection compared with MLN-primed cells from the same donors.

## DISCUSSION

An open question in mucosal immunology is how identical microbial antigens can elicit qualitatively different adaptive immune responses depending on anatomical context ^4^. Using commensal SFB as a model for microbiota-induced T-cell differentiation, we show that PPs and MLNs generate transcriptionally, clonally, and functionally distinct T-cell responses driven by SFB colonization. Rather than inducing uniform Th17 immunity across gut-associated lymphoid tissues, SFB triggers compartmentalized responses in which the site of priming licenses effector capacity, cell trafficking, and ultimately protective potential.

Within this framework, LysoDCs emerge as specialized luminal commensal samplers that establish a unique Th17-priming niche in PPs. Our combined imaging, *ex vivo* priming, and genetic depletion data support a model in which CCR2-dependent Embigin⁺ LysoDCs in the subepithelial dome capture luminal SFB, migrate to interfollicular regions, and selectively prime naive SFB-reactive CD4 T cells under steady-state conditions. Although *in vivo* depletion strategies affect other mononuclear phagocyte populations in addition to LysoDCs, our *ex vivo* co-culture experiments provide the most direct evidence for this role. Since conventional DCs retain full capacity to present exogenous SFB peptide, the selective ability of Embigin⁺ LysoDCs to prime from endogenously acquired antigen points to their non-redundant function as luminal antigen samplers in PPs rather than a general superiority in antigen presentation. This positions LysoDCs as a key APC subset distinct from the CCR2-independent populations implicated in SFB-driven Th17 activation in MLNs ^17,21,30^. While the path to SFB-Th17 activation in MLNs appears to involve a multi-step APC network between intestinal macrophages, migratory cDC2s, and lamina propria-resident cells ^17,20,30,34^, our data support a more unified process in PPs, where a single cell type can both sample luminal antigen and directly prime Th17 cells. This is facilitated by the strategic positioning of LysoDCs in the SED, which provides easy access to luminal commensal antigens, and a short migration pathway into adjacent interfollicular regions to engage naive T cells. Together, these findings reinforce PPs as a parallel site capable of initiating robust effector responses during physiological SFB colonization.

Single-cell analyses further reveal that PP priming generates a qualitatively different T-cell response compared to MLN priming. PP-derived cells upregulate coordinated activation, metabolic adaptation, and tissue-residency modules, with PP-derived Th17 cells already expressing Tr1-like transcriptional features at the priming stage, which suggests that this SFB-induced regulatory program can be initiated locally within PPs rather than exclusively at the lamina propria effector site ^34^. Clonal analysis reinforces this compartmentalized picture, revealing preferential expansion and plasticity within Th17-Tfh17 lineages in PPs. Notably, a recent study showed that SFB colonization can drive Th17 cells in PPs to transdifferentiate into atypical Tfh cells with marked migratory capacity toward the spleen, where they promote extraintestinal autoimmunity ^39^. While our expanded clones are compatible with such plasticity, the tissue-retention signatures we observe in PP-primed cells contrast with the reported migratory phenotype of these atypical cells, a discrepancy that warrants further investigation. More generally, the mechanisms driving local retention of activated T cells in PPs and how these differ from the recirculation-associated programs adopted by MLN-primed cells remain to be resolved.

Nevertheless, regardless of PPs quantitative contribution to the lamina propria pool of SFB-reactive T cells under homeostasis, our functional data demonstrate that PP and MLN priming generate distinct programs with direct consequences for mucosal immunity. In an adoptive transfer model into *C. rodentium*-infected *Rag1*-deficient recipients, PP-primed T cells preserved barrier integrity and ameliorated colitis-associated pathology, whereas MLN-primed cells did not confer comparable protection. Although homeostatic expansion in lymphopenic hosts may amplify pre-existing differences between the transferred populations, both groups showed comparable CD4 T-cell reconstitution in the colonic lamina propria, indicating that the differences in disease outcome observed between groups are unlikely to result from disparities in tissue homing and instead support a durable role for site-specific priming in shaping the functional potential of CD4⁺ T cells.

In summary, we identify PP LysoDCs as a specialized interface between luminal SFB and the adaptive immune system, extending the SFB model beyond simple Th17 induction. More fundamentally, our findings demonstrate that anatomical context shapes the functional identity of commensal-reactive T cells from the earliest stages of priming, with lasting consequences for intestinal homeostasis and mucosal defense.

### Limitations of the study

While our data establish that LysoDCs are required for efficient SFB-specific priming in PPs, the mechanisms underlying their unique priming capacity remain unclear, including whether specific cytokines, co-stimulatory molecules, or metabolite transfer are involved. Similarly, the molecular basis of SFB-specific priming in MLNs requires further investigation, particularly regarding the relative contributions of lamina propria-derived migratory cDC2s and local APC networks. Our functional transfer model in *Rag1*-deficient mice reveals that PP and MLN priming imprint CD4 T cells with distinct protective capacities. However, the use of lymphopenic recipients does not allow assessment of whether and how these two T-cell pools interact and cooperate during physiological pathogen clearance in immunocompetent hosts. Competitive transfer experiments or compartment-specific ablation strategies in intact mice would be needed to address this question. Additionally, determining the ability of PP-primed T cells to migrate from PPs to the lamina propria, as well as their longevity and recall capacity, will be important to fully define their contribution to intestinal immune homeostasis. Finally, although our study relies on murine SFB for mechanistic clarity, the recent demonstration that SFB are prevalent members of the human gut microbiota suggests that analogous PP APC-T cell circuits may operate in humans. Defining these circuits will be essential for translational relevance.

## Supporting information

Resource table

## RESOURCE AVAILABILITY

### Lead contact

Further information and requests for reagents may be directed to and will be fulfilled by the lead contact, Valérie Gaboriau-Routhiau.

## Materials availability

This study did not generate new unique reagents.

## Data and code availability

● Sequencing data have been deposited at NCBI BioProject database (http://www.ncbi.nlm.nih.gov/bioproject/) and are publicly available as of the date of publication. Accession number will be listed in the key resources table.
● This paper does not report original code. References to all code used are available in the STAR Methods section.
● Any additional information required to reanalyze the data reported in this paper is available from the lead contact upon request.

## ACKNOWLEDGMENTS

We thank Dr. Olivier Disson (Pasteur Institute, Paris) for generously providing a key mouse line, Dr. Nicolas Serafini (Pasteur Institute, Paris) for sharing *Citrobacter rodentium* strain, and Marie-Claire Blache (CIML, Marseille) for helpful discussion on image segmentation. We are grateful to the LEAT Animal Husbandry (mainly Amaury Gensou and Emilie Panafieu) and Single Cell Platform (Mélodie Perin, Francesco Carbone, and Maria Emilia Puig Lombardi) at the Imagine Institute for technical support, as well as to the histology platform at INEM and PICSL imaging facility of the CIML (ImagImm), a member of the national infrastructure France-BioImaging and supported by the French National Research Agency (ANR-24-INBS-0005 FBI - BIOGEN), Région SUD, IBiSA and Amidex foundation. This work was supported by institutional funding from INSERM, INRAe, and CNRS, and by grants from the Agence Nationale de la Recherche (ANR-20-CE15-0016, ANR-23-CE15-0018-01 and ANR-24-CE15-2556), the INSERM Transversal Program on Microbiota, and the Fondation Princesse Grace de Monaco. The Imagine Institute is supported by the Investissement d’Avenir program (ANR-10-IAHU-01).

## AUTHOR CONTRIBUTION

Conceptualization, R.O.C., M.C., H.L., and V.G.R.; Methodology, R.O.C., M.C., H.L., and V.G.R.;

Investigation, R.O.C., M.C., M.M.R., A.R., R.H., C.L., H.L., and V.G.R.; Formal Analysis, R.O.C., R.H., C.L., M.F., R.G.A.G., H.L.; Writing – Original Draft, R.O.C.; Writing – Review & Editing, all the authors; Funding Acquisition, H.L., and V.G.R.; Resources, B.J., N.C.B., H.L., and V.G.R.; Supervision, B.J., N.C.B., H.L., and V.G.R.

## DECLARATION OF INTERESTS

The authors declare no competing interests.

## INCLUSION AND DIVERSITY

We support inclusive, diverse, and equitable conduct of research.

## DECLARATION OF AI AND AI-ASSISTED TECHNOLOGIES IN THE WRITING PROCESS

During the preparation of this work, the authors used Claude (Anthropic) and ChatGPT (OpenAI) for writing assistance, text editing, and structural feedback. The authors reviewed and edited the output as needed and take full responsibility for the content of the published article.

## STAR METHODS

### Key Resource Tables

See section below.

## EXPERIMENTAL MODEL AND SUBJECT DETAILS

### Mice

Specific-pathogen free (SPF) wild-type, 7B8 transgenic, LyzM-GFP ^40^, *Ccr2*^-/-^ ^41^ (kindly provided by Dr. Olivier Disson, Pasteur Institute, Paris), germ-free wild-type, and germ-free *Rag1^-/-^* (all C57BL/6 background) were bred at the facilities of IMAGINE Institute. SPF mice were maintained in ventilated cages, while germ-free, SFB-monocolonized, and SPF SFB-negative 7B8 mice were maintained in plastic isolators. All mice had free access to autoclaved water and commercial diet (R03-40; UAR), the latter sterilized by γ-irradiation (40 kGy) for mice kept in isolators. All experiments were performed using age and gender matched groups. All animal procedures were performed according to European guidelines, being previously approved by the French governmental authorities (ethics protocol #9132).

### SFB monoassociation and quantification

SFB monoassociation was achieved by two different methods, as detailed in the main text and/or figure legend. First, SFB were vertically transmitted to the pups by ex-germ-free SFB-monoassociated breeding pairs previously colonized with frozen stocks of a single batch of SFB-enriched feces. Second, adult germ-free mice were oral gavaged for two consecutive days, using 0.3 mL of cecal content supernatant from ex-germ-free SFB-monoassociated mice (also previously monocolonized as the described breeding pairs). This second method was also used to enhance SFB levels at weaning time (postnatal day 21) in SPF mice when needed, as for *Ccr2*^-/-^ mice experiments. Quantification of SFB was performed by extracting bacterial DNA from ileum biopsies with a ribolyzer FastPrep® and Lysing Matrix B beads (MP Biomedical) (adapted from ^42^). RT-PCR was done on a CFX384 TM Real-time system (Bio-Rad) using 16S rDNA primer pairs specific for SFB (see Key Resource Table) and SYBR-Green PCR master mix. SFB levels were quantified using comparative threshold cycle method (2^-ΔCt^) with a standard curve generated with quantified plasmid DNA where SFB 16S rDNA gene had been cloned in PCR-blunt II vector (ThermoFisher) according to suppliers’ instructions.

### Immune cells extraction from PPs, MLNs, and villus LP

After euthanasia, MLNs and small intestine were harvested. PPs were removed from small intestine, cut into small pieces and digested into 6 mL of RPMI containing type VIII Collagenase (100 U/mL) (Sigma) and 175 U/mL of DNase I (Sigma) for 15 min at 37 °C. MLNs were smashed in 70 μm cell strainers and passing cells were collected in cold PBS containing 2% fetal bovine serum. Small intestines were opened longitudinally, washed in cold PBS to remove fecal content, cut into 1 cm pieces, and incubated in 3 mM PBS-EDTA solution for 10 min at 37 °C under agitation. Supernatant was removed and incubation was repeated 3-4 times until a clean supernatant was obtained. Villus lamina propria (LP) immune cells were isolated by digesting the tissue pieces in RPMI with 20% fetal bovine serum containing type VIII Collagenase (100 U/mL) and DNase I (175 U/mL) for 40 min at 37 °C under agitation, followed by mechanical dissociation. Cells were further purified using a Percoll gradient (80:40). For all samples, cells were counted using LUNA (Logos Biosystems) and immediately used for flow cytometry staining.

### Flow cytometry and cell sorting

In 96-well round-bottom plates, cells were first incubated for 5 min at room temperature with 2.4G2 antibody to block Fc receptors, followed by surface markers staining for 20 min at room temperature. LIVE/DEAD™ Fixable Aqua Dead Cell Stain Kit (Life Technologies) was used to evaluate cell mortality. When needed, cells were also stained with Streptavidin conjugated antibodies for 5 min at room temperature, fixed with Foxp3/Transcription Factor Staining Buffer Set (ThermoFisher) for 50 min at room temperature, and finally stained for intracellular markers for 40 min at room temperature. Multiparametric flow cytometry was performed using a LSRFortessa™ X-20 (BD Biosciences) or NovoCyte Penteon (Agilent Technologies), cell sorting was done using FACSAria III (BD Biosciences), and data analysis was performed using FlowJo^TM^ software version 10 (BD Biosciences).

### Naive T cell adoptive transfers

Germ-free mice were monocolonized with SFB by oral gavage as described above. Naive 7B8 CD4 T cells were purified from spleen of 7B8.CD45.1 transgenic SPF mice using EasySep™ Mouse Naive CD4^+^ T Cell Isolation Kit (STEMCELL Technologies), labelled with CellTrace™ Violet (CTV, ThermoFisher) and transferred intravenously to SFB-monocolonized (5x10^5^ cells/mouse), seven days after gavage. Recipient mice were euthanized 24, 48, or 72 hours (h) after transfer. Similar experiment was performed using SPF SFB-negative (CD45.2) mice as recipients (euthanized 72 h after transfer). PPs and MLNs were isolated and processed as described above.

### FTY720 immunosuppressive treatment

Germ-free mice were monocolonized with SFB by oral gavage, as mentioned above. Mice were injected intraperitoneally with 1 mg/kg FTY720 (Sigma-Aldrich) for five or seven consecutive days, with the first injection done immediately after the first SFB gavage. Control mice were injected with daily doses of 0.1% sodium chloride. At day five or seven after colonization, mice were euthanized and PPs and MLNs were harvested for flow cytometry analysis. Peripheral blood samples were analyzed using IDEXX ProCyte DX (IDEXX Laboratories) to confirm systemic lymphopenia and therefore, the efficacy of the treatment.

### Treatment with clodronate-loaded liposomes

Germ-free mice were injected intravenously with 250 µL of clodronate-loaded liposomes or control PBS-liposomes from Liposoma BV (Amsterdam, Netherlands). Five days later, mice were monocolonized with SFB by oral gavage, concomitant with adoptive transfer of naive 7B8 CD45.1 CD4 T cells, both procedures as described above. After five more days, mice were euthanized and PPs and MLNs were harvested for flow cytometry analysis.

### Co-culture assays

Distinct populations of phagocytes were isolated from PPs, MLNs, and villus LP of either SFB-monocolonized or SPF SFB-negative mice, as described above. After FACS sorting, cells were cultured in Cerottini medium (DMEM + Glutamax, L-asparagine (36 µg/mL), L-arginine (116 µg/mL), folic acid (10 µg/mL), HEPES (1 mg/mL), β-mercaptoethanol (10 µM) and 10% fetal bovine serum), containing M-CSF (20 ng/mL) and GM-CSF (20 ng/mL), for 2 hours at 37 °C and 5% CO_2_, in the presence or absence of purified SFB-peptide A6 (2.5 µM) ^29^. In the meantime, splenic naive 7B8 CD4^+^ T cells were purified from 7B8.CD45.1 transgenic SPF mice using EasySep™ Mouse Naive CD4^+^ T Cell Isolation Kit (STEMCELL Technologies) and labelled with CellTrace™ Violet (CTV, ThermoFisher). Naive 7B8 CD4^+^ T cells were then resuspended in Cerottini medium containing IL-2 and IL-7 (both at 5 µg/mL) and co-cultured with the distinct phagocyte subsets at a 1:5 ratio. After 4 days at 37 °C and 5% CO_2_, cells were collected and processed for flow cytometry analysis, as described above.

### Single-cell RNA-sequencing

Germ-free mice were monocolonized with SFB by oral gavage, as described above. Seven days after gavage, mice were euthanized, PPs and MLNs were harvested, and immune cells were isolated and stained for flow cytometry. Cells from individual animals were then hashed using TotalSeq™ anti-mouse Hashtag Antibodies (BioLegend), according to suppliers’ instructions. Single cell suspensions were isolated from a pool of 4 mice for each tissue (PPs and MLNs). Primed (CD44^high^CD62L^-^) CD4^+^ T cells were sorted among live CD45^+^ singlets. The libraries were generated using Chromium Next GEM Single Cell 5’ Reagent Kit v2 with Feature Barcode technology for Cell Surface Protein & Immune Receptor Mapping (10X Genomics) according to the manufacturer’s protocol. Gene expression (mRNA), VDJ TCR and Cell Surface Protein libraries were constructed. Briefly, cells were counted and up to 20,000 cells were loaded in the 10X Genomics Chromium X to generate single-cell gel-beads in emulsion. After reverse transcription, gel-beads in emulsion were disrupted. Barcoded complementary DNA was isolated and amplified by PCR. Following fragmentation, end repair and A-tailing, sample indexes were added during index PCR. The purified libraries were sequenced on a Novaseq 6000 (Illumina) with 26 cycles of read 1, 10 cycles of i7 index, 10 cycles of i5 index and 90 cycles of read 2, targeting a median depth of 40,000 reads per cell for gene expression, 5,000 reads per cell for TCR VDJ library and 5,000 reads per cell for Cell Surface Protein library. Sequencing reads were demultiplexed and aligned to the mouse reference transcriptome (mm10-2020-A), using the CellRanger Pipeline (v8.0.0). The unfiltered RNA UMI counts were loaded into Seurat v5.01 for quality control, data integration and downstream analyses. Cells with a low number of detected genes, a low counts depth and a high fraction of mitochondrial content were considered outliers and removed if they differ by 4 times their median absolute deviation (MAD). Doublets were identified and removed using DoubletFinder and an expected doublet rate of 0.075. Data from each sample were normalized and scaled using the sctransform method. After filtering, the cells were demultiplexed using the HTODemux, and only the cells with unique hashes were assigned back to their sample of origin. Batch effect was corrected using Seurat’s FindIntegratedAnchors, and the samples were integrated using the rpca reduction.

### Single-cell dimensional reduction, clustering, and annotation

On the integrated dataset, dimensional reduction was performed using principal component analysis, and UMAP visualizations were generated using the first 30 principal components. Graph-based clustering was performed in Seurat using the shared nearest-neighbor graph, and the resolution selected for annotation was resolution = 1. Cell identities were annotated using a combined automated and marker-based strategy. First, SingleR was applied to normalized RNA expression data using the ImmGen reference dataset, generating both fine-and main-level immune cell labels with Wilcoxon-based marker detection. In parallel, curated marker sets representing major immune lineages and CD4⁺ T cell states were scored with Seurat’s AddModuleScore function. Cluster-level module scores, SingleR predictions, canonical marker-gene expression, and manual inspection of UMAP, feature, violin, and heatmap visualizations were then integrated to assign final biological identities. The final annotation resolved 19 cell states. After annotation and quality control, 27,380 cells assigned to biologically interpretable clusters were retained for downstream analyses. UMAP visualizations were generated using the final annotated object and colored either by anatomical compartment, defined as mesenteric lymph nodes (MLN) or Peyer’s patches (PP), or by final cluster identity. For organ-stratified visualization, cells from MLN and PP were displayed separately using the same UMAP coordinates and a fixed cluster color scheme to preserve interpretability across panels.

### Compositional analysis of cell-state abundance

To assess whether the cellular composition differed between MLN and PP, cell counts were aggregated at the biological-sample level for each annotated cluster. Each sample was required to map uniquely to a single organ, and samples with fewer than 20 total cells were removed. Clusters were retained for compositional testing only if they contained at least 10 cells in total and were represented in at least two biological samples. Differential abundance of cell states was modeled using sccomp, with sample-level cluster counts as input. The compositional model included organ as the main covariate for both the abundance and variability components: composition ∼ organ and variability ∼ organ. The model was fitted with bimodal mean–variability association enabled and tested using sccomp_test. Organ-associated compositional effects were extracted from the organ term, including posterior effect estimates, credible intervals, hypothesis probabilities, and false-discovery-rate-adjusted composition and variability statistics. Observed sample-level proportions were plotted as violin plots with individual biological samples overlaid, and significant compositional effects were summarized. For unsupervised visualization of sample-level compositional similarity, the sample-by-cluster proportion matrix was zero-adjusted, renormalized, transformed by centered log-ratio transformation, and clustered using Euclidean distance with Ward’s minimum-variance linkage (ward.D2). The resulting hierarchical tree was displayed alongside stacked barplots of cluster proportions for each sample, preserving the dendrogram-derived sample order.

### Pseudobulk differential expression by cluster

Differential gene expression between PP and MLN was performed independently within each annotated cluster using a pseudobulk strategy. For each cluster, raw RNA counts were aggregated by biological sample and organ using the sum of counts across cells. Pseudobulk profiles were retained only when they contained at least 20 cells, and clusters were analyzed only when at least two pseudobulk replicates were available for each organ. Pseudobulk profiles were also computed per biological replicate using individual mouse hashing, ensuring that organ effects were estimated across independent biological samples rather than pooled cells. Mitochondrial genes were removed before differential testing. Lowly expressed genes were filtered using edgeR::filterByExpr, and library sizes were normalized using trimmed mean of M-values normalization. Mean-variance relationships were modeled using limma-voom, followed by linear modeling with empirical Bayes moderation. The design matrix modeled organ as the explanatory variable: ∼ organ with MLN used as the reference level and the PP-vs-MLN contrast extracted from the organPP coefficient. Genes were considered differentially expressed when they showed a Benjamini-Hochberg-adjusted p value < 0.05 and an absolute log₂ fold change > 1. Positive log₂ fold changes were interpreted as higher expression in PP, whereas negative log₂ fold changes were interpreted as higher expression in MLN. Cluster-level differential expression results were visualized using a consolidated dot plot showing significant PP- or MLN-associated genes across clusters.

### Pseudobulk analysis of curated transcriptional modules

To evaluate transcriptional programs associated with Th17, regulatory, activation, TCR signaling, NF-κB tuning, metabolic adaptation, and microbiota responsiveness, raw counts were aggregated into pseudobulk profiles by biological sample within each target cell state. Counts were normalized using TMM normalization in edgeR, and log₂ counts per million were computed with a prior count of 2. For Th17-focused analyses, curated Tr1-like and SFB-responsive gene signatures were evaluated. Genes were matched to the expression matrix in a case-insensitive manner, and missing genes were recorded for transparency. Signature expression matrices were extracted from the pseudobulk logCPM matrix and scaled by gene using row-wise z scores prior to visualization. Heatmaps were generated with pheatmap, using Euclidean distance and Ward.D2 linkage for row and column clustering. For broader module-level analyses across cell types, predefined gene modules were evaluated in the same pseudobulk framework. For each cell type, module genes present in the expression matrix were extracted from the pseudobulk logCPM matrix, scaled by gene, and visualized as heatmaps.

### Glycolysis-associated trajectory analysis

To investigate whether glycolytic transcriptional activity varied along a Th17-Tfh17-related continuum, trajectory analysis was performed on a subset of cells annotated as Th17, Cycling Tfh17, or Tfh17. The Hallmark Glycolysis gene set was obtained from the mouse MSigDB collection using msigdbr, and genes present in the Seurat object were retained. Cells were reprocessed using glycolysis genes as the feature space: the selected genes were scaled, PCA was computed using up to 20 principal components, and UMAP was generated from the glycolysis-based PCA embedding using up to the first 10 PCs. A per-cell glycolysis score was calculated as the mean log-normalized expression of the retained glycolysis genes. The glycolysis-based embedding was converted to a SingleCellExperiment object, and lineage inference was performed using slingshot on the glycolysis UMAP representation. Cluster labels were provided from the final annotation, with Th17 set as the starting cluster and Tfh17 as the terminal cluster. Pseudotime values were extracted for all inferred lineages, and a global pseudotime score was defined as the mean pseudotime across non-missing lineage assignments for each cell. Organ-associated differences in pseudotime and glycolysis score were tested within each cluster using Wilcoxon rank-sum tests, followed by Benjamini-Hochberg correction. To assess whether the relationship between glycolytic activity and pseudotime differed by organ, cluster-stratified linear models were fitted using glycolysis_score ∼ pseudotime_global * organ.

### TCR repertoire analysis

TCR repertoire analyses were performed using the Seurat object containing V(D)J metadata. VDJ metadata were first harmonized by converting logical and numeric fields to consistent formats and by extracting productive TRA and TRB CDR3 sequences where available. Only cells with productive TCR information and a valid overlap identifier were retained for repertoire analyses. The primary overlap definition used a stringent paired-chain key combining the first detected productive TRA and TRB CDR3 sequences (tcr_key_tra_trb). A TRB-only key was also supported by the analysis framework, but the figures listed here were generated using the paired TRA/TRB key. Clone sizes were computed in a sample-aware manner by counting the number of cells sharing the same overlap key within each biological sample. Clones were categorized into expansion classes according to clone size: singleton, small, medium, large, and hyperexpanded. These clone-class annotations were used to quantify clonal expansion by organ and T cell cluster and to generate stacked barplots of clone-class composition. Organ specificity of TCRs was evaluated by classifying each unique TCR key as PP-specific, MLN-specific, or shared between PP and MLN. Global organ-specificity summaries were generated from unique TCR identifiers, whereas cluster-level analyses quantified the number and proportion of shared and organ-restricted TCRs within each T cell state. TCR overlap between organs and between cell-state sectors was quantified using both Jaccard overlap, based on the presence or absence of shared TCR identifiers, and Morisita-Horn similarity, based on abundance-weighted TCR count distributions. Morisita-Horn values were bounded between 0 and 1 for plotting. Pairwise Morisita-Horn similarity between T cell clusters was computed using the paired TRA/TRB overlap key. Heatmaps were generated separately for all organs combined, MLN only, and PP only. Prior to heatmap generation, Cycling Tfh, Tfh2, Naive/Tcm Gilz⁺, and Recently Activated tTreg clusters were excluded from the heatmap universe. Missing or non-finite Morisita-Horn values were set to zero for clustering, diagonal values were set to one, and hierarchical clustering was performed on a distance matrix defined as 1 minus Morisita-Horn similarity using complete linkage. Heatmap color scales were constrained from 0 to 1. To visualize shared TCR architecture across organ-cluster sectors, circular plots were generated using circlize. Sectors were defined by organ and T cell cluster. The outer track represented organ identity, the middle track represented cluster identity, and the inner track represented clone-class composition for singleton, small, and medium clones. Links between sectors represented shared TCR identifiers and were anchored to the corresponding clone-class segment within each sector. Links connecting sectors within the same organ were colored using the organ color, whereas links connecting MLN and PP sectors were colored as shared TCRs. TCR V-gene usage was summarized by extracting the first detected TRB V gene from each productive TCR record and calculating gene frequencies by cluster and organ. Organ-associated differences in TRB V-gene frequency were represented as PP-minus-MLN frequency differences and visualized across clusters using fixed organ-aware color scales. Additional TRA and TRB V/J-pair visualizations were generated to summarize global and organ-stratified V–J pairing structure, using fixed gene-color assignments to preserve comparability across panels.

### Citrobacter rodentium infection

A single colony of the bioluminescent *Citrobacter rodentium* strain ICC180 was cultured overnight in Luria-Bertani (LB) broth supplemented with kanamycin (40 µg/mL). The culture was subsequently diluted and grown in fresh LB containing kanamycin to mid-log phase. Bacteria were then pelleted by centrifugation and resuspended in sterile PBS. Germ-free *Rag1*-deficient mice were orally gavaged with 2×10⁹ colony-forming units (CFU) in 200 µL. Body weight was monitored daily, and experiments were terminated when weight loss exceeded 20% of the initial body weight. Bacterial burden was later determined by weighing feces, liver, and spleen samples, homogenizing them in PBS, and plating serial dilutions onto LB agar supplemented with kanamycin for CFU enumeration. 24 hours post-infection, lymphopenic mice received intravenously either PBS or primed CD4 T cells sorted from PP or MLN of SFB monocolonized mice, prepared as described above.

### Immunofluorescence staining and confocal microscopy

Biological samples were isolated and fixed with Antigenfix (Diapath) for 1 hour at 4 °C, washed and either processed for paraffin inclusion or incubated in 30% sucrose overnight, followed by OCT embedding, freezing and storage at -20 °C. For paraffin samples, sections were cut at 4 μm thickness and stained with hematoxylin-eosin or Periodic acid-Schiff. For OCT, sections were cut at 20 μm thickness and stored at -20 °C. After permeabilization and blocking of unspecific binding sites for 30 min with PBS containing 0.5% saponin, 1% fetal bovine serum, 2% BSA, and 1% donkey serum, sections were labelled with primary antibodies diluted in the same blocking buffer overnight at 4 °C, washed in PBS and incubated with secondary antibodies at room temperature for 1 hour. Then, sections were washed in PBS and incubated with serum from the primary antibody species for 30 minutes, followed by a 1-hour staining in the same blocking solution using antibodies conjugated to fluorochromes. Slides were mounted in Prolong Gold (Thermofisher) and visualized with a Zeiss LSM 980 confocal microscope equipped with spectral imaging mode ^43^. Images were analyzed with Zeiss Zen lite, QuPath ^44^, Cellpose ^45^, Imaris 10.1 (Oxford Instruments) and Adobe Photoshop CC 2022.

### Quantification and statistical analysis

Most data are presented as box-and-whisker plots, unless otherwise stated in the figure legends. In box-and-whisker plots, the center line indicates the median, the box indicates the interquartile range, and whiskers indicate the minimum and maximum values. Data were assessed for normality using the Kolmogorov-Smirnov test and analyzed accordingly using either an unpaired two-tailed t-test or, when appropriate, the Mann-Whitney U test. For comparisons involving multiple groups or variables, one-way or two-way ANOVA followed by Sidak’s or Tukey’s multiple-comparisons test was used, as indicated in the figure legends. A p-value < 0.05 was considered statistically significant in all analyses, with *p < 0.05, **p < 0.01, ***p < 0.001, and ****p < 0.0001 displayed on the figures. Statistical analyses were performed using GraphPad Prism version 10 for Windows (GraphPad Software).

---

**Supplementary Figure 1.**
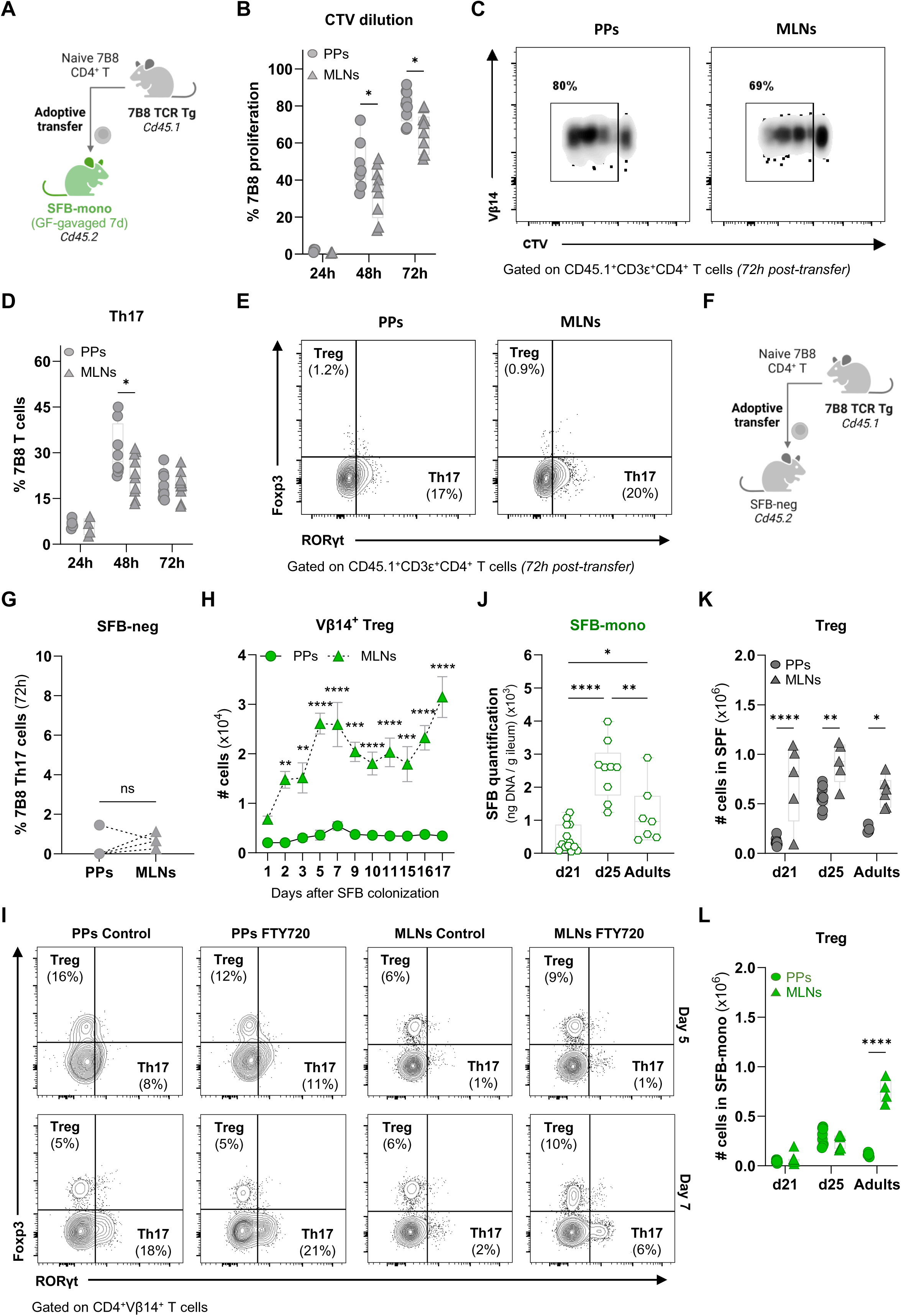
: (A) Schematic of adoptive transfer of naive 7B8 CD45.1 CD4 T cells from SPF mice into adult ex-germ-free SFB-monoassociated CD45.2 mice colonized with SFB by oral gavage seven days prior to transfer. Analyses were performed 24, 48, and 72 h after adoptive transfer (related to B-E). (B) Frequency of 7B8 T cell proliferation based on CTV^TM^ dilution in PPs (circles) and MLNs (triangles) over time. Data show individual values for each mouse in box-and-whisker plots. Two independent experiments, n = 4-10 mice/group. Statistical analysis by two-way ANOVA with Sidak’s multiple-comparisons test. (C) Representative Vâ14/CTV FACS plots, gated on live CD45.1^+^CD3ε^+^CD4^+^ T cells, from PPs and MLNs, 72 h post-transfer. (D) Frequency of 7B8 T cell differentiation into Th17 cells in PPs (circles) and MLNs (triangles) over time. Two independent experiments, n = 4-10 mice/group. Statistical analysis by two-way ANOVA with Sidak’s multiple-comparisons test. (E) Representative Foxp3/RORγt FACS plots, gated on live CD45.1^+^CD3ε^+^CD4^+^ T cells, 72h post-transfer. (F) Schematic of adoptive transfer of naive 7B8 CD45.1 CD4 T cells from SPF mice to SPF SFB-negative CD45.2 mice (related to G). (G) Frequency of 7B8 T cell differentiation into Th17 cells in PPs (circles) and MLNs (triangles) 72 h post transfer. Data show individual values for each mouse in box-and-whisker plots. One experiment, n = 4 mice. Statistical analysis by paired two-tailed t-test. ns = not significant. (H) Numbers of SFB-specific Treg (Foxp3^+^RORγt^-^) cells in PPs (circles) and MLNs (triangles) in SFB-monocolonized mice over time. Each symbol represents mean ± SEM. Two independent experiments, n = 4-10 mice/group. Statistical analysis by two-way ANOVA with Sidak’s multiple-comparisons test. (I) Representative Foxp3/RORãt FACS plots, gated on live CD4^+^ Vβ14^+^ T cells, from PPs and MLNs of both control and FTY720-treated mice. (J) SFB quantification in the terminal ileum of SFB-monocolonized mice, naturally colonized from birth, by qPCR with primers specific to SFB 16S rDNA gene at different time points: postnatal days 21 (d21), 25 (d25) or 45-60 (adults). Data show individual values for each mouse in box-and-whisker plots. Three independent experiments, n = 7-13 mice. Statistical analysis by one-way ANOVA with Tukey’s multiple-comparisons test. (K-L) Numbers of Treg (Foxp3^+^RORγt^-^) cells from SPF mice (K) and SFB-monocolonized mice (L) in PPs (circles) or MLNs (triangles) at the indicated time points. Data show individual values for each mouse in box-and-whisker plots. Three independent experiments, n = 4-9 mice/group. Statistical analysis by two-way ANOVA with Sidak’s multiple-comparisons test.

**Supplementary Figure 2 :**
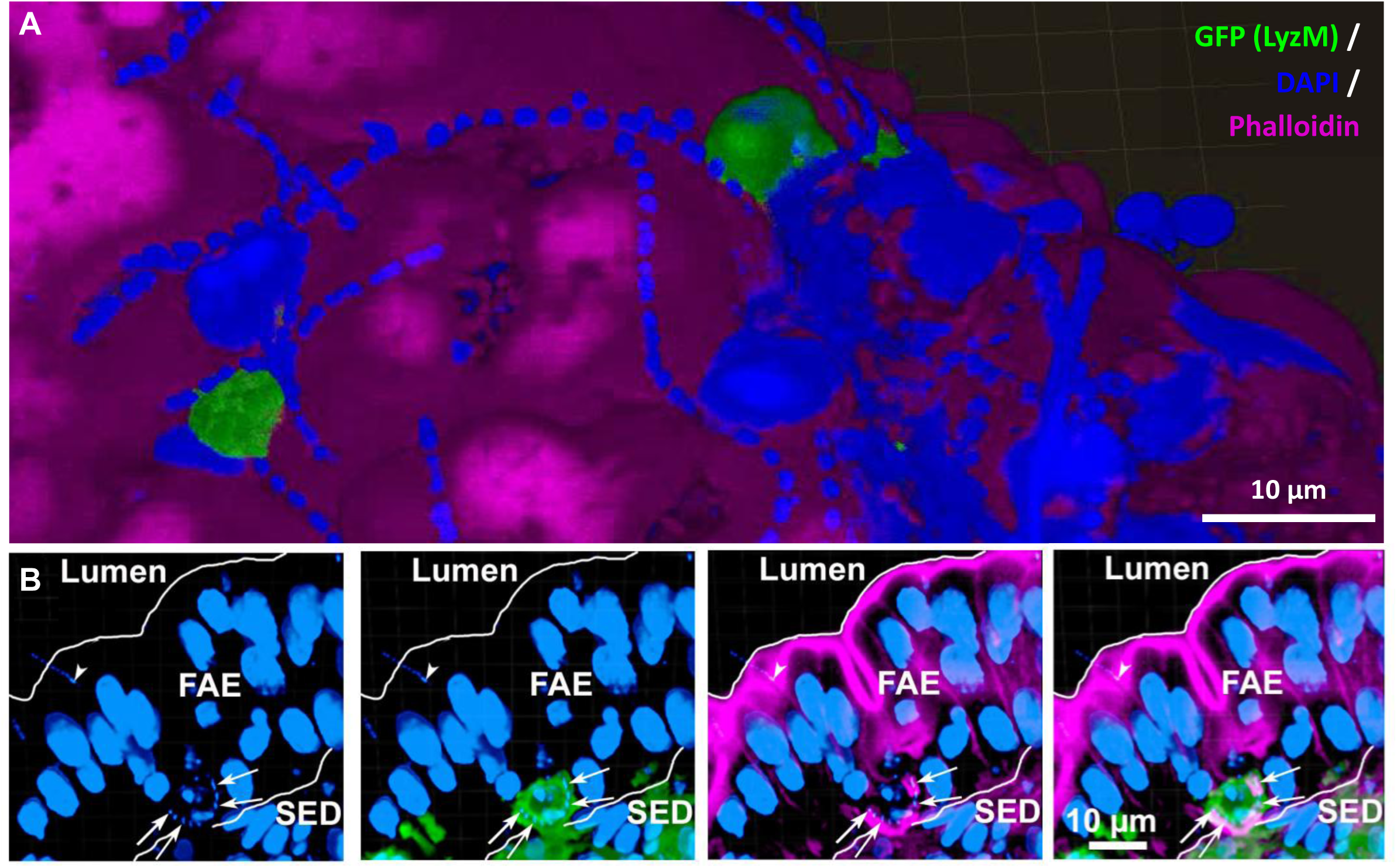
(A) Surface view of a PP from a lysozyme M (LyzM)-GFP mouse showing two LysoDC trans-M cell dendrites (GFP, green) extending through the follicle-associated epithelium (FAE, phalloidin, magenta) into the lumen, in close contact with SFB (small DAPI-stained dot chains, cyan). (B) Section of a PP from a LyzM-GFP mouse showing an SFB (cyan dots, arrows) internalized by a LysoDC (green). The internalized SFB is surrounded by a dense filamentous actin network (magenta). An arrowhead marks another SFB anchored in the FAE. SED = subepithelial dome.

**Supplementary Figure 3 :**
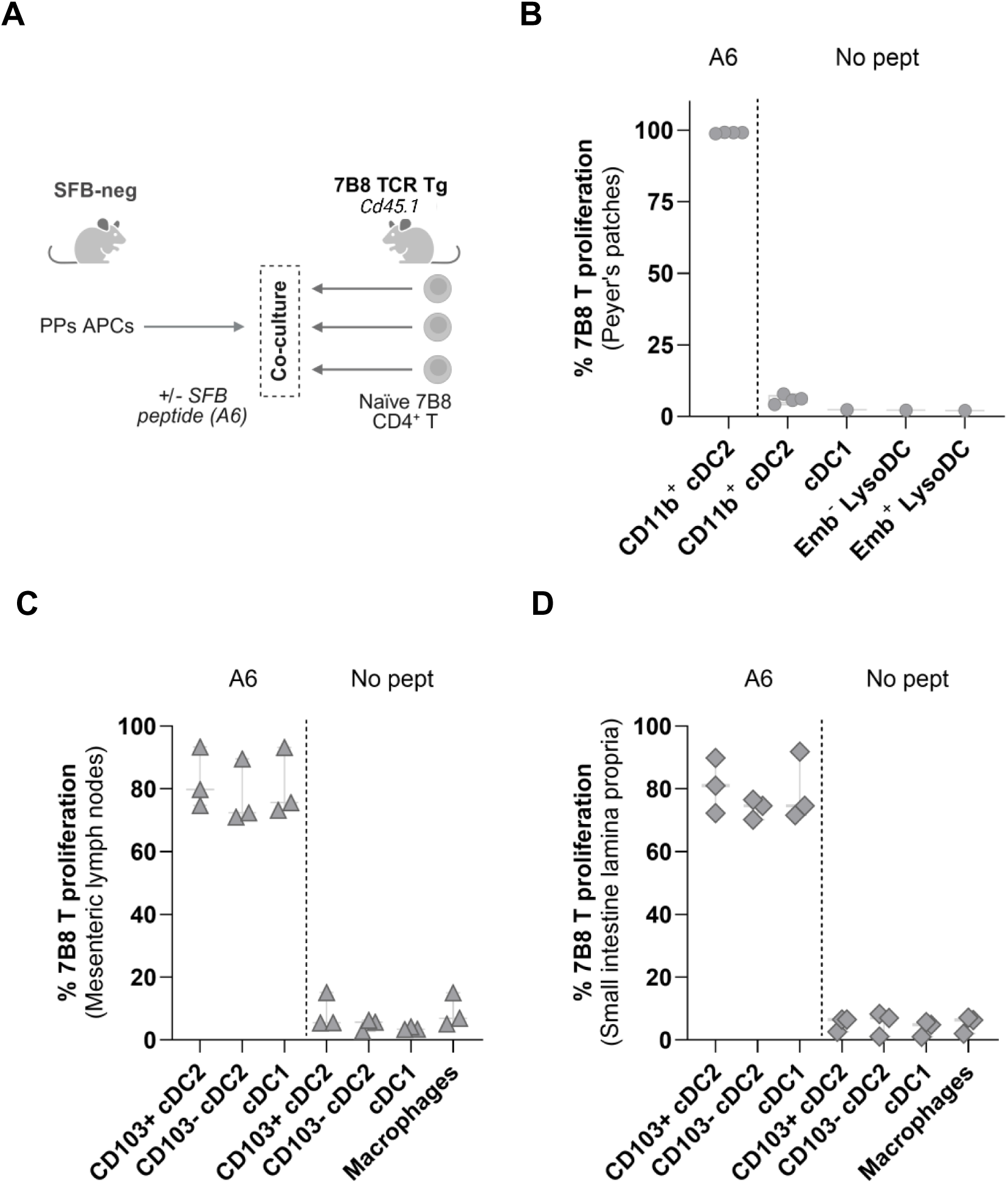
(A) Scheme of co-culture of distinct FACS-sorted APCs from PPs of SPF SFB-negative mice with naive 7B8 CD4 T cells, with or without SFB-specific A6 peptide. Cell proliferation was assessed by CTV^TM^ dilution after four days in culture (related to B). (B) Frequency of 7B8 T cell proliferation after co-culture. One experiment. APCs were FACS-sorted from pooled PPs of 15-20 mice, and 1-4 replicate co-cultures were established per condition. (C-D) Frequency of 7B8 T cell proliferation after co-culture with MLN APCs (C) or small intestine lamina propria APCs (D) in the presence or absence of exogenous A6 peptide. Three independent experiments. In each experiment, APCs were FACS-sorted from pooled tissues of 15-20 mice, and 2-5 replicate co-cultures were established per condition. (B-D) Statistical analysis by one-way ANOVA with Sidak’s multiple-comparisons test.

**Supplementary Figure 4 :**
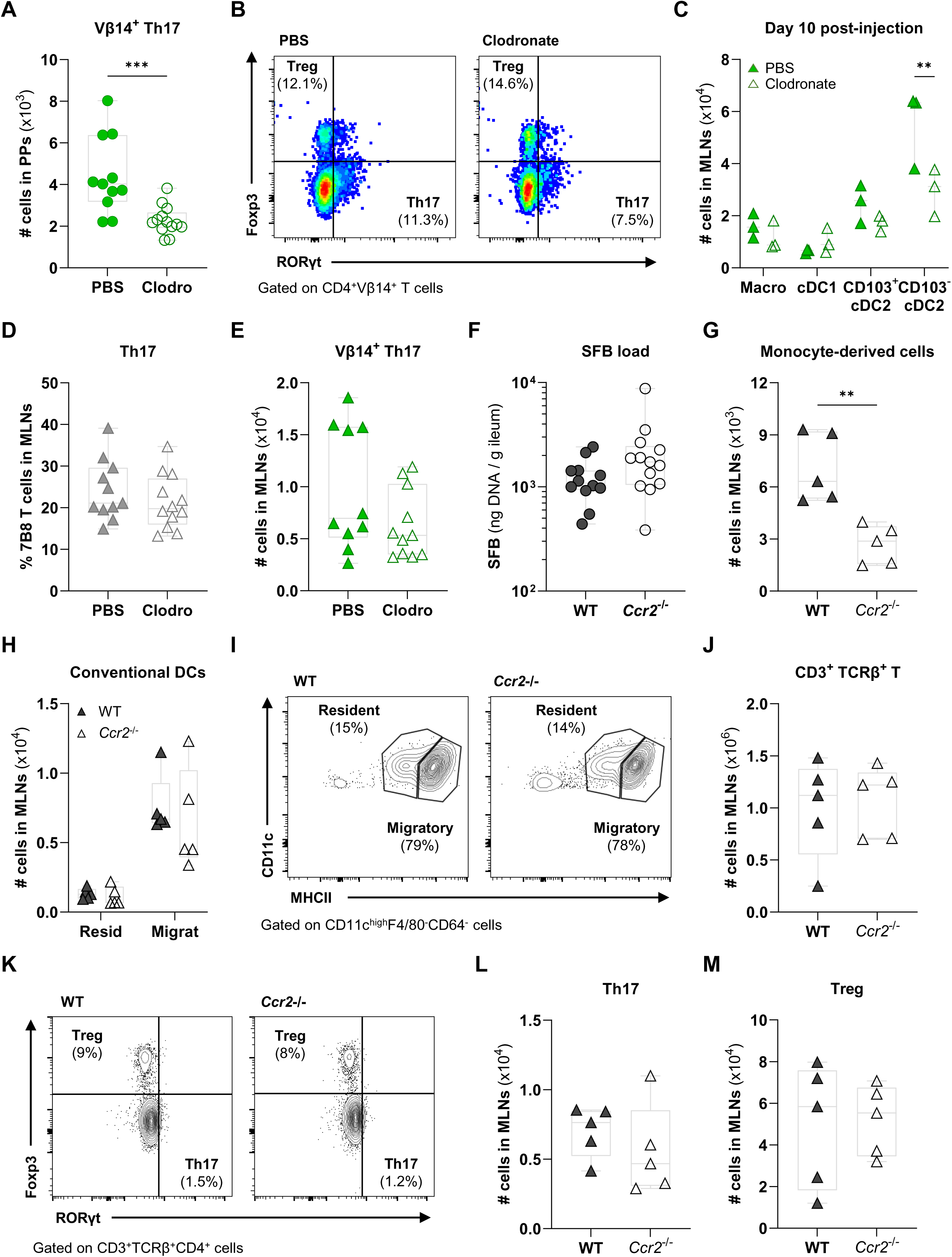
(A) Numbers of endogenous Vâ14^+^ Th17 (Foxp3^-^RORγt^+^) polarized cells in PPs of clodronate-treated mice (empty symbols) or PBS-control mice (filled symbols) at day 10 post-liposome injection. (B) Representative Foxp3/RORγt FACS plots gated on live CD4^+^ Vβ14^+^ T lymphocytes in PPs of each group. (C) Number of mononuclear phagocytes in mesenteric lymph nodes (MLNs) of each group. (D-E) Percentages of 7B8 T cells (D) and numbers of endogenous Vâ14^+^ T cells (E) polarized towards Th17 (FoxP3^-^RORγt^+^) cells in MLNs of each group. (A-E) Data show individual values for each mouse in box-and-whisker plots. Two independent experiments, n = 10-13 mice/group. Statistical analysis by unpaired two-tailed t-test (A, D-E), or by two-way ANOVA with Sidak’s multiple-comparisons test (C). (F-M) Comparisons between wild-type (WT, filled symbols) and *Ccr2^-/-^* mice (empty symbols). For quantified panels, data show individual values for each mouse in box-and-whisker plots. Two independent experiments, n = 6-8 mice/group. Statistical analysis by unpaired two-tailed t-test (F-G, J, L-M), or by two-way ANOVA with Sidak’s multiple-comparisons test (H). (F) SFB quantification in the terminal ileum by qPCR with primers specific to SFB 16S rDNA gene. (G-H) Number of monocyte-derived cells (G), and resident and migratory conventional dendritic cells (H) in MLNs. (I) Representative CD11c/MHCII FACS plots gated on live CD11c^high^ F4/80^-^ CD64^-^ cells in MLNs. (J) Number of CD3ε^+^ TCRβ^+^ T cells in MLNs. (K) Representative Foxp3/RORγt FACS plots, gated on live CD3ε^+^ TCRβ^+^ CD4^+^ cells in MLNs. (L-M) Number of Th17 cells (L), and Treg cells (M) in MLNs.

**Supplementary Figure 5 :**
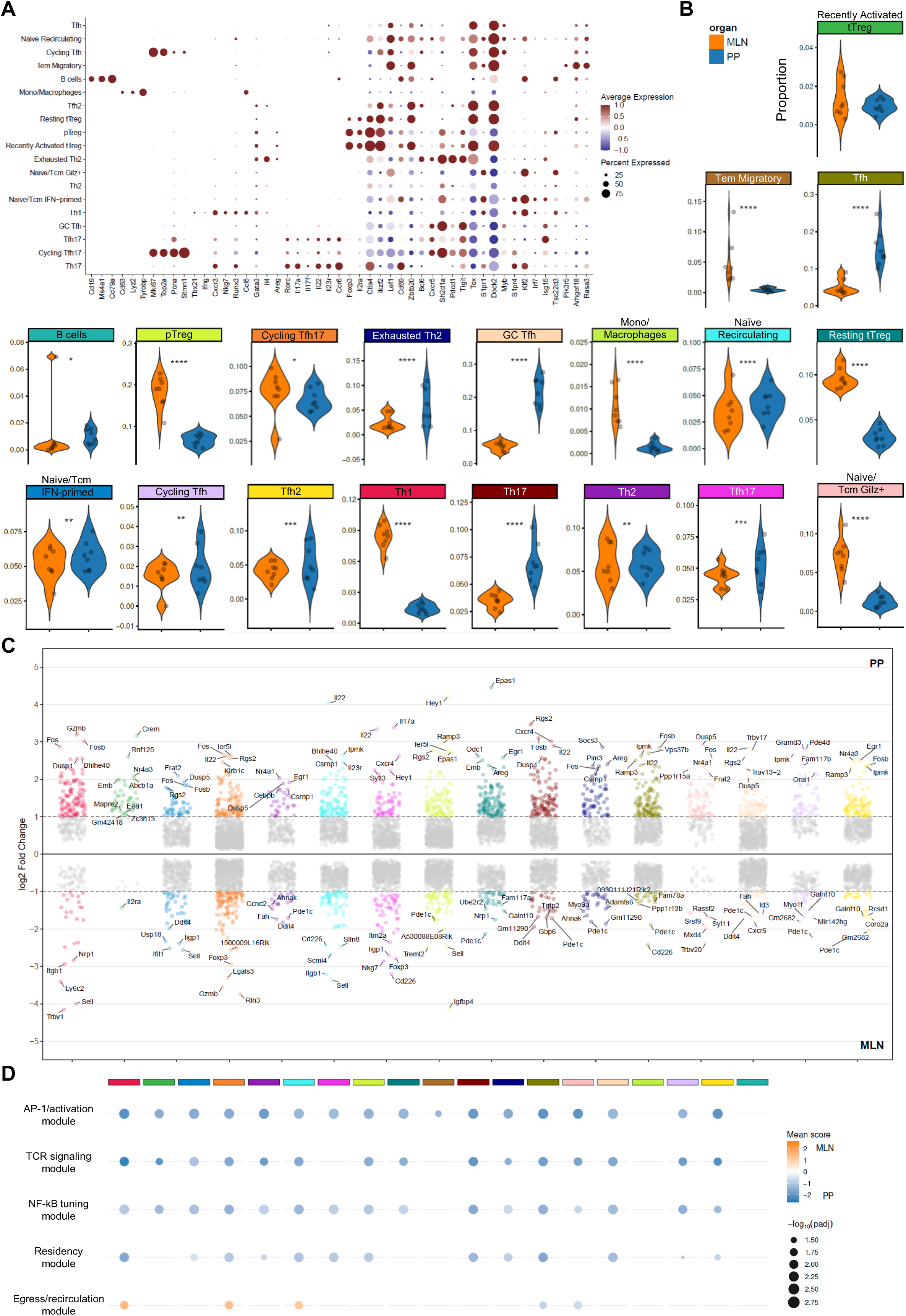
(A) Dot plot showing expression of selected marker genes across all 19 annotated clusters. Dot size indicates the percentage of cells expressing the gene; color intensity indicates average expression level (z-scored). (B) Violin plots showing the proportion of each individual cluster in MLN (orange) versus PP (blue). Statistical analysis by sccomp compositional modeling (see STAR Methods). (C) Differential gene expression between PP and MLN across all annotated clusters. Each column represents a cluster, and each dot represents a differentially expressed gene. Positive log₂ fold change (top, toward PP) and negative log₂ fold change (bottom, toward MLN) are shown. The top five most significantly differentially expressed genes for each tissue are highlighted. Differential expression by pseudobulk limma-voom with Benjamini-Hochberg correction (adjusted p < 0.05, |log₂FC| > 1). (D) Dot plot summarizing module score differences between PP and MLN across all annotated clusters. Dot size indicates −log₁₀ (adjusted p-value); color indicates mean score difference (blue = PP-enriched, orange = MLN-enriched). Module score comparisons by Wilcoxon rank-sum test with Benjamini-Hochberg correction.

**Supplementary Figure 6 :**
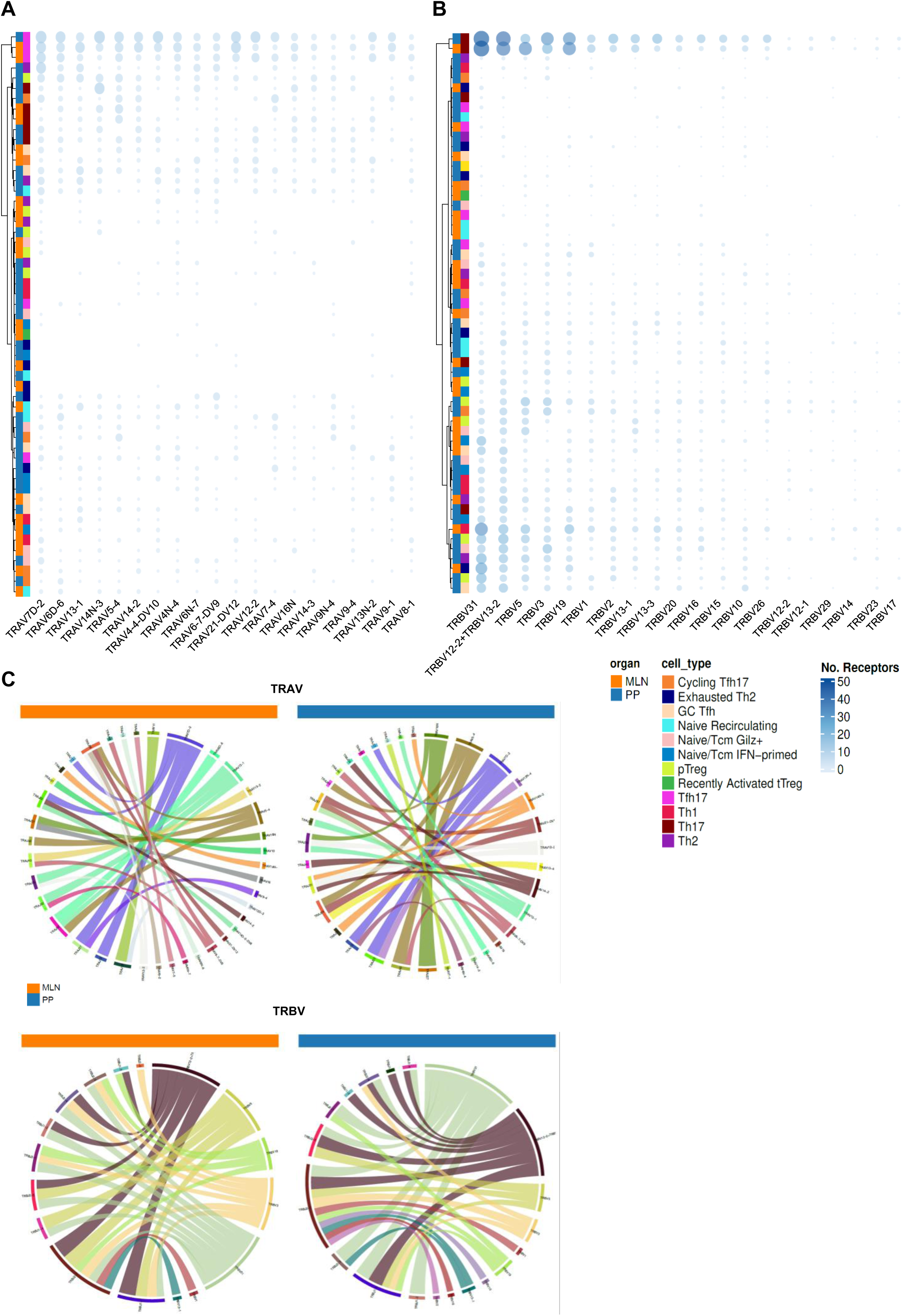
(A) TRAV gene usage across clonotypes, stratified by tissue origin (PP, blue; MLN, orange) and cell type (color-coded by cluster identity as in Figure 5A). Each column represents a TRAV gene segment; rows represent individual cell clusters from PP or MLN. Dot size indicates the number of receptors using each gene segment. (B) TRBV gene usage, displayed in the same format as (A). (C) Chord diagrams showing V-gene pairing patterns stratified by tissue. Left two panels: TRAV gene usage in MLN (orange) and PPs (blue). Right two panels: TRBV gene usage in MLN (orange) and PPs (blue). Arc width is proportional to the frequency of each V-gene pairing. Colors correspond to cell type identity as in Figure 5A.

**Supplementary Figure 7 :**
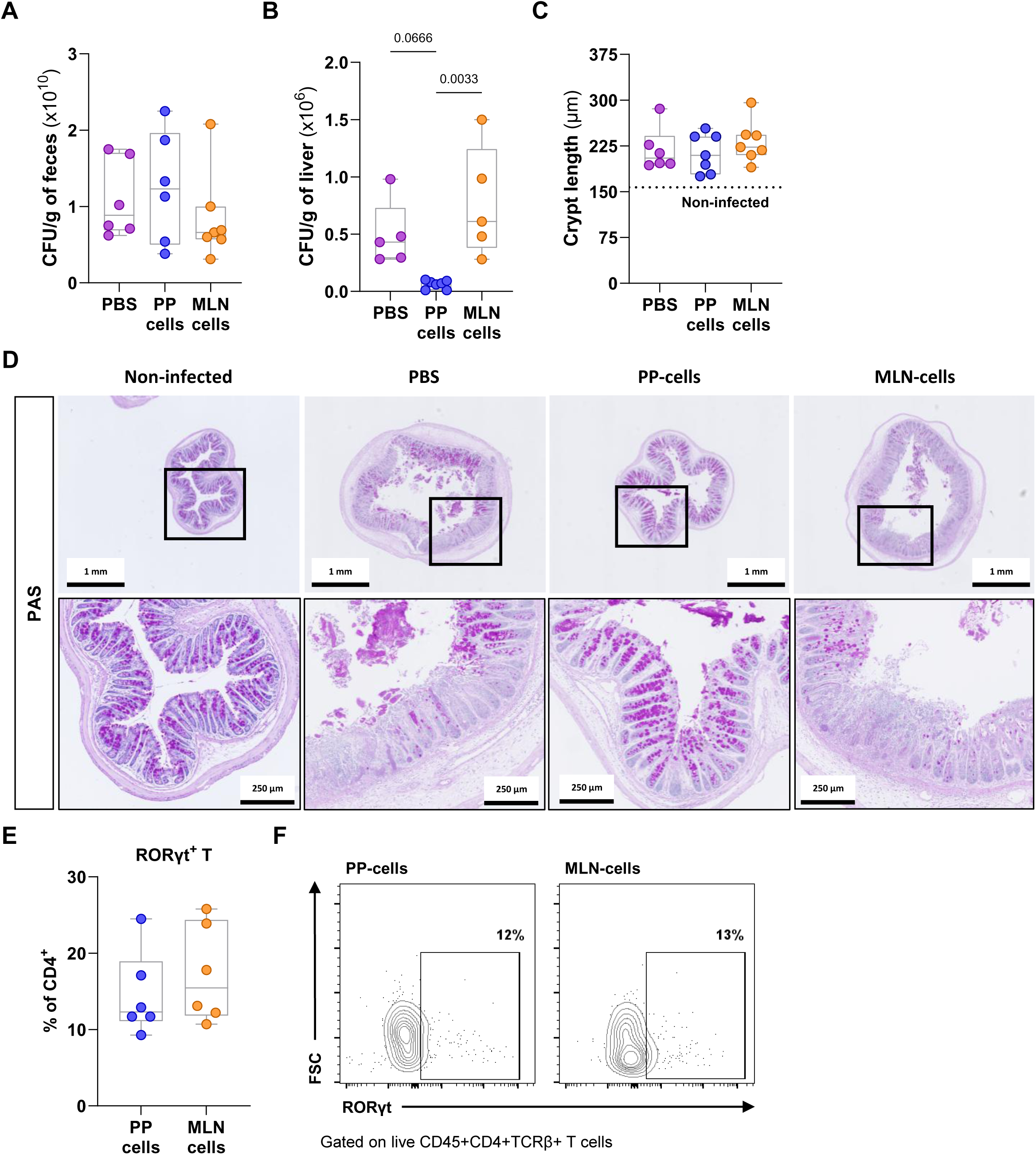
(A-B) CR bacterial burden in the feces (A) or in the liver (B), quantified by culture and CFU counting, and normalized to the sample weight. Data show individual values for each mouse in box-and-whisker plots. (C) Crypt length (µm) measured from H&E-stained sections. The dotted line indicates the mean crypt length from non-infected germ-free *Rag1*-deficient mice. (D) Representative PAS-stained colon sections from *Rag1*-deficient mice, either non-infected or infected with CR and receiving PBS or the indicated primed cells. (E) Frequency of RORãt⁺ T cells among CD4⁺TCRβ⁺ T cells from the colonic lamina propria. Data show individual values in box-and-whisker plots. Statistical analysis by unpaired two-tailed t-test. (F) Representative RORγt/FSC FACS plots, gated on live CD45⁺CD4⁺TCRβ⁺ T cells.

## Supplementary Movie S1 (related to Figure 2)

3D reconstruction of an interfollicular LysoDC (green) in close interaction (contact zones in blue) with two proliferating (Ki67 in yellow) SFB-specific CD45.1^+^ T cells (magenta) in PPs, 72 hours after transfer into a CD45.2^+^ LyzM-GFP mouse.

## Supplementary Movie S2 (related to Figure 2)

3D reconstruction of two interfollicular LysoDCs (green) in close interaction (contact zones in blue) with two SFB-specific CD45.1^+^ T cells (magenta), one of which has differentiated into a Th17 cell (RORγt^+^ in orange, Foxp3^-^ in red) in PPs, 72 hours after transfer into a CD45.2^+^ LyzM-GFP mouse.

## Notes

### Competing Interest Statement

The authors have declared no competing interest.

## REFERENCES

1. Gensollen, T., Iyer, S.S., Kasper, D.L., and Blumberg, R.S. (2016). How colonization by microbiota in early life shapes the immune system. Science 352, 539–544. 10.1126/science.aad9378.

2. Million, M., Tomas, J., Wagner, C., Lelouard, H., Raoult, D., and Gorvel, J.-P. (2018). New insights in gut microbiota and mucosal immunity of the small intestine. Hum. Microb. J. 7-8, 23–32. 10.1016/j.humic.2018.01.004.

3. Al Nabhani, Z., Dulauroy, S., Marques, R., Cousu, C., Al Bounny, S., Déjardin, F., Sparwasser, T., Bérard, M., Cerf-Bensussan, N., and Eberl, G. (2019). A Weaning Reaction to Microbiota Is Required for Resistance to Immunopathologies in the Adult. Immunity 50, 1276–1288.e5. 10.1016/j.immuni.2019.02.014.

4. Kedmi, R., and Littman, D.R. (2024). Antigen-presenting cells as specialized drivers of intestinal T cell functions. Immunity 57, 2269–2279. 10.1016/j.immuni.2024.09.011.

5. Mayassi, T., Barreiro, L.B., Rossjohn, J., and Jabri, B. (2021). A multilayered immune system through the lens of unconventional T cells. Nature 595, 501–510. 10.1038/s41586-021-03578-0.

6. Canesso, M.C.C., Castro, T.B.R., Nakandakari-Higa, S., Lockhart, A., Luehr, J., Bortolatto, J., Parsa, R., Esterházy, D., Lyu, M., Liu, T.-T., et al. (2024). Identification of antigen-presenting cell–T cell interactions driving immune responses to food. Science (1979). 10.1126/science.ado5088.

7. Kiran, S., Cruz, A.R., Daniau, A., Ma, B., Marbouty, M., Pipoli Da Fonseca, J., Legrand, A., Baudry, L., Cokelaer, T., Bensussan, M., et al. (2026). Segmented filamentous bacteria are worldwide human gut commensals. Nat. Commun. 17. 10.1038/s41467-026-70010-4.

8. Gaboriau-Routhiau, V., Rakotobe, S., Lécuyer, E., Mulder, I., Lan, A., Bridonneau, C., Rochet, V., Pisi, A., De Paepe, M., Brandi, G., et al. (2009). The key role of segmented filamentous bacteria in the coordinated maturation of gut helper T cell responses. Immunity 31, 677–689. 10.1016/j.immuni.2009.08.020.

9. Ivanov, I.I., Atarashi, K., Manel, N., Brodie, E.L., Shima, T., Karaoz, U., Wei, D., Goldfarb, K.C., Santee, C.A., Lynch, S. V, et al. (2009). Induction of intestinal Th17 cells by segmented filamentous bacteria. Cell 139, 485–498. 10.1016/j.cell.2009.09.033.

10. Lécuyer, E., Rakotobe, S., Lengliné-Garnier, H., Lebreton, C., Picard, M., Juste, C., Fritzen, R., Eberl, G., McCoy, K.D., Macpherson, A.J., et al. (2014). Segmented filamentous bacterium uses secondary and tertiary lymphoid tissues to induce gut IgA and specific T helper 17 cell responses. Immunity 40, 608–620. 10.1016/j.immuni.2014.03.009.

11. Chung, H., Pamp, S.J., Hill, J.A., Surana, N.K., Edelman, S.M., Troy, E.B., Reading, N.C., Villablanca, E.J., Wang, S., Mora, J.R., et al. (2012). Gut immune maturation depends on colonization with a host-specific microbiota. Cell 149, 1578–1593. 10.1016/j.cell.2012.04.037.

12. Burgess, S.L., Buonomo, E., Carey, M., Cowardin, C., Naylor, C., Noor, Z., Wills-Karp, M., and Petri, W.A. (2014). Bone marrow dendritic cells from mice with an altered microbiota provide interleukin 17A-dependent protection against Entamoeba histolytica colitis. mBio 5, e01817. 10.1128/mBio.01817-14.

13. McAleer, J.P., Nguyen, N.L.H., Chen, K., Kumar, P., Ricks, D.M., Binnie, M., Armentrout, R.A., Pociask, D.A., Hein, A., Yu, A., et al. (2016). Pulmonary Th17 Antifungal Immunity Is Regulated by the Gut Microbiome. J. Immunol. 197, 97–107. 10.4049/jimmunol.1502566.

14. Shi, Z., Zou, J., Zhang, Z., Zhao, X., Noriega, J., Zhang, B., Zhao, C., Ingle, H., Bittinger, K., Mattei, L.M., et al. (2019). Segmented Filamentous Bacteria Prevent and Cure Rotavirus Infection. Cell 179, 644–658.e13. 10.1016/j.cell.2019.09.028.

15. Lai, N.Y., Musser, M.A., Pinho-Ribeiro, F.A., Baral, P., Jacobson, A., Ma, P., Potts, D.E., Chen, Z., Paik, D., Soualhi, S., et al. (2020). Gut-Innervating Nociceptor Neurons Regulate Peyer’s Patch Microfold Cells and SFB Levels to Mediate Salmonella Host Defense. Cell 180, 33–49.e22. 10.1016/j.cell.2019.11.014.

16. Woo, V., Eshleman, E.M., Hashimoto-Hill, S., Whitt, J., Wu, S.-E., Engleman, L., Rice, T., Karns, R., Qualls, J.E., Haslam, D.B., et al. (2021). Commensal segmented filamentous bacteria-derived retinoic acid primes host defense to intestinal infection. Cell Host Microbe 29, 1744–1756.e5. 10.1016/j.chom.2021.09.010.

17. Sano, T., Kageyama, T., Fang, V., Kedmi, R., Martinez, C.S., Talbot, J., Chen, A., Cabrera, I., Gorshko, O., Kurakake, R., et al. (2021). Redundant cytokine requirement for intestinal microbiota-induced Th17 cell differentiation in draining lymph nodes. Cell Rep. 36, 109608. 10.1016/j.celrep.2021.109608.

18. Welty, N.E., Staley, C., Ghilardi, N., Sadowsky, M.J., Igyártó, B.Z., and Kaplan, D.H. (2013). Intestinal lamina propria dendritic cells maintain T cell homeostasis but do not affect commensalism. Journal of Experimental Medicine 210, 2011–2024. 10.1084/jem.20130728.

19. Goto, Y., Panea, C., Nakato, G., Cebula, A., Lee, C., Diez, M.G., Laufer, T.M., Ignatowicz, L., and Ivanov, I.I. (2014). Segmented filamentous bacteria antigens presented by intestinal dendritic cells drive mucosal Th17 cell differentiation. Immunity 40, 594–607. 10.1016/j.immuni.2014.03.005.

20. Panea, C., Farkas, A.M., Goto, Y., Abdollahi-Roodsaz, S., Lee, C., Koscsó, B., Gowda, K., Hohl, T.M., Bogunovic, M., and Ivanov, I.I. (2015). Intestinal Monocyte-Derived Macrophages Control Commensal-Specific Th17 Responses. Cell Rep. 12, 1314–1324. 10.1016/j.celrep.2015.07.040.

21. Ngoi, S., Yang, Y., Iwanowycz, S., Gutierrez, J., Li, Y., Williams, C., Hill, M., Chung, D., Allen, C., and Liu, B. (2022). Migrating Type 2 Dendritic Cells Prime Mucosal Th17 Cells Specific to Small Intestinal Commensal Bacteria. The Journal of Immunology 209, 1200–1211. 10.4049/jimmunol.2200204.

22. Morikawa, M., Tsujibe, S., Kiyoshima-Shibata, J., Watanabe, Y., Kato-Nagaoka, N., Shida, K., and Matsumoto, S. (2016). Microbiota of the Small Intestine Is Selectively Engulfed by Phagocytes of the Lamina Propria and Peyer’s Patches. PLoS One 11, e0163607. 10.1371/journal.pone.0163607.

23. Da Silva, C., Wagner, C., Bonnardel, J., Gorvel, J.-P., and Lelouard, H. (2017). The Peyer’s Patch Mononuclear Phagocyte System at Steady State and during Infection. Front. Immunol. 8, 1254. 10.3389/fimmu.2017.01254.

24. Luciani, C., Hager, F.T., Cerovic, V., and Lelouard, H. (2022). Dendritic cell functions in the inductive and effector sites of intestinal immunity. Mucosal Immunol. 15, 40–50. 10.1038/s41385-021-00448-w.

25. Lelouard, H., Fallet, M., de Bovis, B., Méresse, S., and Gorvel, J.-P. (2012). Peyer’s patch dendritic cells sample antigens by extending dendrites through M cell-specific transcellular pores. Gastroenterology 142, 592–601.e3. 10.1053/j.gastro.2011.11.039.

26. Wagner, C., Bonnardel, J., Da Silva, C., Spinelli, L., Portilla, C.A., Tomas, J., Lagier, M., Chasson, L., Masse, M., Dalod, M., et al. (2020). Differentiation Paths of Peyer’s Patch LysoDCs Are Linked to Sampling Site Positioning, Migration, and T Cell Priming. Cell Rep. 31, 107479. 10.1016/j.celrep.2020.03.043.

27. Bonnardel, J., Da Silva, C., Henri, S., Tamoutounour, S., Chasson, L., Montañana-Sanchis, F., Gorvel, J.-P., and Lelouard, H. (2015). Innate and adaptive immune functions of peyer’s patch monocyte-derived cells. Cell Rep. 11, 770–784. 10.1016/j.celrep.2015.03.067.

28. Martínez-López, M., Iborra, S., Conde-Garrosa, R., Mastrangelo, A., Danne, C., Mann, E.R., Reid, D.M., Gaboriau-Routhiau, V., Chaparro, M., Lorenzo, M.P., et al. (2019). Microbiota Sensing by Mincle-Syk Axis in Dendritic Cells Regulates Interleukin-17 and -22 Production and Promotes Intestinal Barrier Integrity. Immunity 50, 446–461.e9. 10.1016/j.immuni.2018.12.020.

29. Yang, Y., Torchinsky, M.B., Gobert, M., Xiong, H., Xu, M., Linehan, J.L., Alonzo, F., Ng, C., Chen, A., Lin, X., et al. (2014). Focused specificity of intestinal TH17 cells towards commensal bacterial antigens. Nature 510, 152–156. 10.1038/nature13279.

30. Goto, Y., Panea, C., Nakato, G., Cebula, A., Lee, C., Diez, M.G., Laufer, T.M., Ignatowicz, L., and Ivanov, I.I. (2014). Segmented filamentous bacteria antigens presented by intestinal dendritic cells drive mucosal Th17 cell differentiation. Immunity 40, 594–607. 10.1016/j.immuni.2014.03.005.

31. Jiang, H.Q., Bos, N.A., and Cebra, J.J. (2001). Timing, localization, and persistence of colonization by segmented filamentous bacteria in the neonatal mouse gut depend on immune status of mothers and pups. Infect. Immun. 69, 3611–3617. 10.1128/IAI.69.6.3611-3617.2001.

32. Metwaly, A., Jovic, J., Waldschmitt, N., Khaloian, S., Heimes, H., Häcker, D., Ahmed, M., Hammoudi, N., Le Bourhis, L., Mayorgas, A., et al. (2023). Diet prevents the expansion of segmented filamentous bacteria and ileo-colonic inflammation in a model of Crohn’s disease. Microbiome 11, 66. 10.1186/s40168-023-01508-y.

33. De Giovanni, M., Vykunta, V.S., Biram, A., Chen, K.Y., Taglinao, H., An, J., Sheppard, D., Paidassi, H., and Cyster, J.G. (2024). Mast cells help organize the Peyer’s patch niche for induction of IgA responses. Sci. Immunol. 9, eadj7363. 10.1126/sciimmunol.adj7363.

34. Brockmann, L., Tran, A., Huang, Y., Edwards, M., Ronda, C., Wang, H.H., and Ivanov, I.I. (2023). Intestinal microbiota-specific Th17 cells possess regulatory properties and suppress effector T cells via c-MAF and IL-10. Immunity 56, 2719–2735.e7. 10.1016/j.immuni.2023.11.003.

35. Liu, Y., Xu, D., Guo, S., Wang, S., Ding, H., Siu, C., and Wan, F. (2024). The gut microbiota-independent virulence of noninvasive bacterial pathogen Citrobacter rodentium. PLoS Pathog. 20, e1012758. 10.1371/journal.ppat.1012758.

36. Zindl, C.L., Wilson, C.G., Chadha, A.S., Duck, L.W., Cai, B., Harbour, S.N., Nagaoka-Kamata, Y., Hatton, R.D., Gao, M., Figge, D.A., et al. (2024). Distal colonocytes targeted by C. rodentium recruit T-cell help for barrier defence. Nature 629, 669–678. 10.1038/s41586-024-07288-1.

37. Gudmundsdottir, H., and Turka, L.A. (2001). A closer look at homeostatic proliferation of CD4+ T cells: costimulatory requirements and role in memory formation. J. Immunol. 167, 3699–3707. 10.4049/jimmunol.167.7.3699.

38. Min, B., Yamane, H., Hu-Li, J., and Paul, W.E. (2005). Spontaneous and homeostatic proliferation of CD4 T cells are regulated by different mechanisms. J. Immunol. 174, 6039–6044. 10.4049/jimmunol.174.10.6039.

39. Fan, T., Tai, C., Sleiman, K.C., Cutcliffe, M.P., Kim, H., Liu, Y., Li, J., Xin, G., Grashel, M., Baert, L., et al. (2025). Aberrant T follicular helper cells generated by TH17 cell plasticity in the gut promote extraintestinal autoimmunity. Nat. Immunol. 26, 790–804. 10.1038/s41590-025-02125-7.

40. Faust, N., Varas, F., Kelly, L.M., Heck, S., and Graf, T. (2000). Insertion of enhanced green fluorescent protein into the lysozyme gene creates mice with green fluorescent granulocytes and macrophages. Blood 96, 719–726.

41. Boring, L., Gosling, J., Chensue, S.W., Kunkel, S.L., Farese, R. V, Broxmeyer, H.E., and Charo, I.F. (1997). Impaired monocyte migration and reduced type 1 (Th1) cytokine responses in C-C chemokine receptor 2 knockout mice. Journal of Clinical Investigation 100, 2552–2561. 10.1172/JCI119798.

42. Godon, J.J., Zumstein, E., Dabert, P., Habouzit, F., and Moletta, R. (1997). Molecular microbial diversity of an anaerobic digestor as determined by small-subunit rDNA sequence analysis. Appl. Environ. Microbiol. 63, 2802–2813. 10.1128/aem.63.7.2802-2813.1997.

43. Lelouard, H., Mailfert, S., and Fallet, M. (2018). A Ten-color Spectral Imaging Strategy to Reveal Localization of Gut Immune Cell Subsets.

44. Bankhead, P., Loughrey, M.B., Fernández, J.A., Dombrowski, Y., McArt, D.G., Dunne, P.D., McQuaid, S., Gray, R.T., Murray, L.J., Coleman, H.G., et al. (2017). QuPath: Open source software for digital pathology image analysis. Sci. Rep. 7, 16878. 10.1038/s41598-017-17204-5.

45. Stringer, C., Wang, T., Michaelos, M., and Pachitariu, M. (2021). Cellpose: a generalist algorithm for cellular segmentation. Nat. Methods 18, 100–106. 10.1038/s41592-020-01018-x.

